# *Obox4* promotes zygotic genome activation upon loss of *Dux*

**DOI:** 10.1101/2022.07.04.498763

**Authors:** Youjia Guo, Tomohiro Kitano, Kimiko Inoue, Kensaku Murano, Michiko Hirose, Ten D. Li, Akihiko Sakashita, Hirotsugu Ishizu, Narumi Ogonuki, Shogo Matoba, Masayuki Sato, Atsuo Ogura, Haruhiko Siomi

## Abstract

Once fertilized, mouse zygotes rapidly proceed to zygotic genome activation (ZGA), during which long terminal repeats (LTRs) of murine endogenous retroviruses with leucine tRNA primer (MERVL) are activated by a conserved homeodomain-containing transcription factor, DUX. However, *Dux*-knockout embryos produce fertile mice, suggesting that ZGA is redundantly driven by an unknown factor(s). Here we present multiple lines of evidence that the multicopy homeobox gene, *Obox4*, encodes a transcription factor that is highly expressed in mouse 2-cell embryos and redundantly drives ZGA. Genome-wide profiling revealed that OBOX4 specifically binds and activates MERVL LTRs as well as a subset of murine endogenous retroviruses with lysine tRNA primer (MERVK) LTRs. Depletion of *Obox4* is tolerated by embryogenesis, whereas concomitant *Obox4*/*Dux* depletion markedly compromises embryonic development. Our study identified OBOX4 as a transcription factor that provides genetic redundancy to pre-implantation development.

## Introduction

The mechanism by which zygotes acquire totipotency is a major question in Developmental Biology. Following fertilization, the zygote must build a *de novo* conceptus with a transcriptionally quiescent genome. The genome rapidly undergoes zygotic genome activation (ZGA), during which epigenetic reprogramming and expression of nascent transcripts enforce the replacement of parental infrastructures by their zygotic counterparts ^1^. Completion of metazoan ZGA results in blastomeres, a collection of cells that are totipotent enough to reflect their potential to individually produce both embryos and extraembryonic appendages ^2,3^. Totipotency lasts until the morula stage, where the first cell fate decision takes place, following which it is remolded into pluripotency in cells destined for the embryonic lineage ^4^, to which embryonic stem cells (ESCs) and induced pluripotent stem cells (iPSCs) correspond. In placental mammals, ZGA is characterized by the massive reactivation of transposable elements (TEs) and epigenome remodeling, predominantly implemented by a collection of ZGA genes that have adapted long terminal repeats (LTRs) of endogenous retroviruses (ERVs) as stage-specific *cis*-regulatory elements ^5^.

Homeodomains are DNA-binding amino acid motifs encoded by a class of conserved genomic sequences termed homeoboxes ^6–8^. Homeodomains are prevalent transcription factors that regulate development because of their ability to bind chromatin in a sequence-specific manner ^9–11^. Among all homeobox-containing genes, Paired-like (PRD-like) homeobox class genes are particularly associated with ZGA ^12^. Studies have shown that the PRD-like homeobox gene, double homeobox (*Dux*), plays an important role in pre-implantation embryogenesis ^13,14^. *Dux* is conserved throughout placentalia and is highly expressed during ZGA. In humans and mice, DUX specifically binds and activates endogenous retroviruses with leucine tRNA primer (ERVL) LTR-derived promoters, resulting in the expression of downstream ZGA genes ^15–17^. *Dux* expression induces a 2-cell-embryo-like (2C-like) state in mouse embryonic stem cells (mESCs) ^15–17^, whereas *Dux*-knockout (KO) undermines mouse blastocyst formation ^15^. However, subsequent studies have revealed that mouse embryogenesis is compatible with the loss of *Dux*, suggesting a redundant pathway of ZGA that is controlled by uncharacterized transcription factor(s) ^18–21^.

In this study, we sought to identify the redundant transcription factor that drives ZGA in the absence of *Dux*. We discovered that the multicopy homeobox gene, oocyte-specific homeobox 4 (*Obox4*), promotes *Dux*-less ZGA. *Obox4* is abundantly expressed in mouse 2C-embryos, 2C-like mESCs, and totipotent blastomere-like cells (TBLCs) ^22^. Mechanistically, OBOX4 promotes ZGA by binding to the LTRs of murine endogenous retroviruses with leucine tRNA primer (MERVL) and murine endogenous retroviruses with lysine tRNA primer (MERVK), and thereby affecting the deposition of active epigenetic modifications. Concomitant, but not respective, depletion of *Obox4* and *Dux* severely compromises ZGA and preimplantation development. Taken together, our findings substantiate a pre-implantation development model, in which the ZGA is redundantly promoted by *Dux* and *Obox4*.

## Results

### Expression profiling identifies transcription factor candidates

Genetic redundancy is commonly observed among genes that share sequence homology ^23^. Because *Dux* is a homeobox gene, we speculated that its redundant factors are homeobox genes. We examined the expression profiles of all mouse homeobox genes during pre-implantation embryogenesis using published single-cell RNA-seq (scRNA-seq) data ^24^. Clustering analysis defined a collection of 45 homeobox genes, whose transcript levels reached a maximum before the late 2C stage (Fig. 1a). We then sought to identify *bona fide* ZGA factors from this collection by looking for genes whose transcripts were of zygotic origin, instead of ones that were parentally inherited. TBLCs have recently been established *as in vitro* counterparts of 2C-embryos, which are derived from splicing-inhibited mESCs, and hence, are free of parentally inherited transcripts ^22^. Analysis of published RNA sequencing (RNA-seq) data showed that the expression of four homeobox genes, *Duxf3*, *Emx2*, *Hoxd13*, and *Obox4*, was ten times higher in TBLCs than in ESCs (Fig. 1b). We examined whether the expression of these genes was affected in the *Dux* knockout embryos. Using published RNA-seq data ^18,20^, we found that the expression of all single-copy candidate genes and a subset of the multicopy gene *Obox4* was unaffected in *Dux* knockout embryos (Fig. 1c).

**Figure 1.**
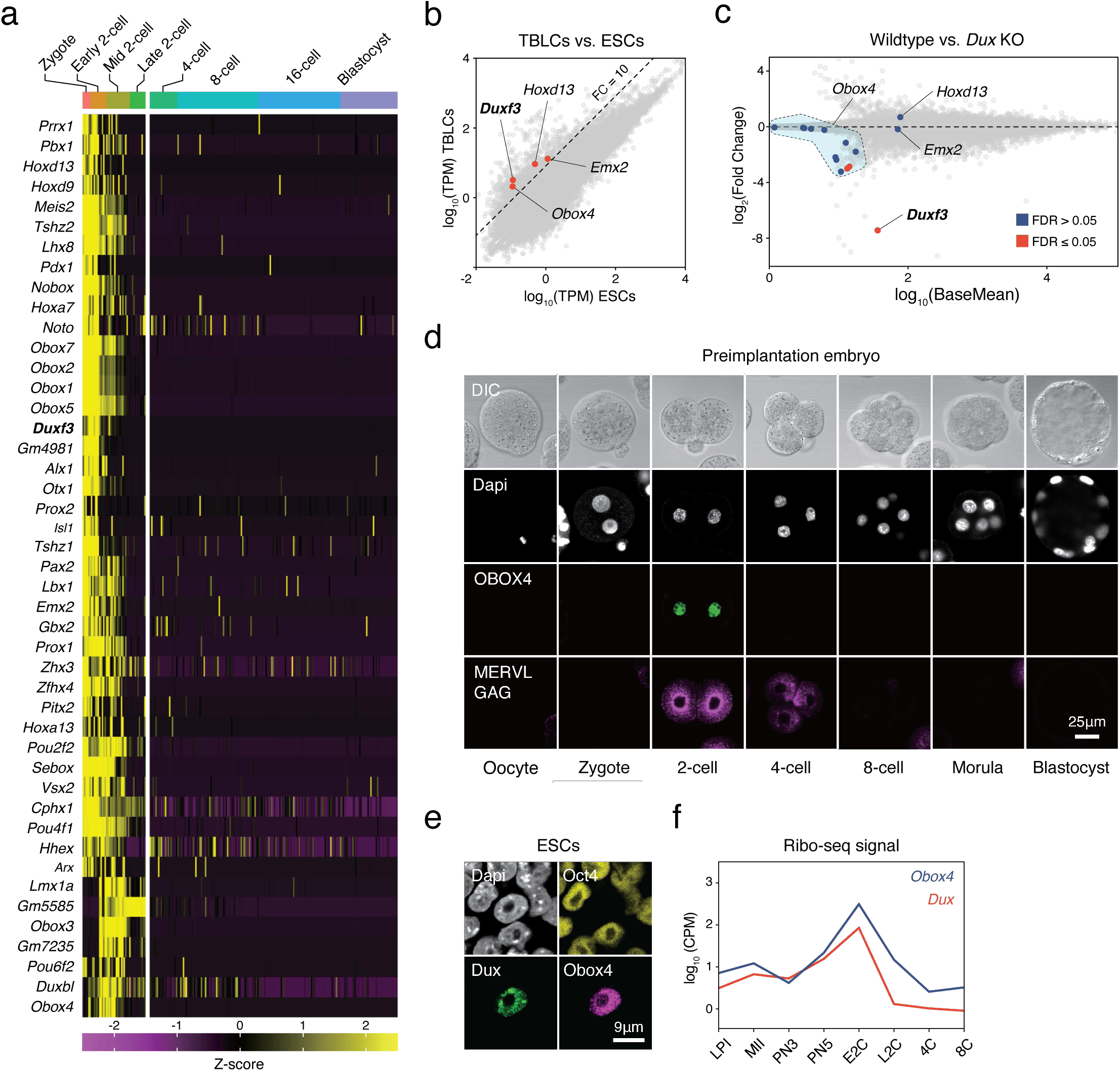
OBOX4 is expressed during ZGA. a) Mouse homeobox genes that are specifically expressed during ZGA. The genes were identified by means of statistically determined *k*-means clustering based on their expression in pre-implantation embryos. *Dux* is shown in a bold italic font. b) Scatterplot showing per-gene normalized read counts in mouse totipotent TBLCs *versus* ESCs. Genes with more than 10-fold normalized read counts (FC=10) have been highlighted. *Dux* is shown in a bold italic font. c) MA plot displaying gene expression in *Dux* knockout 2C-embryos *versus* WT 2C-embryos. Loss of *Dux* partially deactivates the multicopy homeobox gene *Obox4*. d) Immunofluorescence staining of OBOX4 and MERVL GAG at different pre-implantation embryo stages. e) Immunofluorescence staining of DUX, OBOX4, and OCT4 in 2C-like ESCs. f) Translation profile of DUX and OBOX4 during preimplantation characterized by Ribo-seq signal.

### *Obox4* activates 2C-genes and TEs in mESC

Before examining whether the candidates were crucial for embryogenesis, we first confirmed the presence of endogenous OBOX4 protein, as the *Obox4* loci are marked as pseudogenes in the current genome annotation GRCm39^25^. We generated mouse anti-OBOX4 monoclonal antibodies. Immunofluorescence staining using these antibodies confirmed that endogenous OBOX4 was expressed in zygotes and highly abundant in 2C-embryos and 2C-like mESCs (Fig. 1d-e), which is consistent with translatome profile captured by Ribo-seq that detects OBOX4 translation at 2C stage ^26^ (Fig. 1f). Mouse ZGA is characterized by the surging activity of genes that specifically express at 2C stage (2C-gene), many of which have co-opted MERVL LTRs as stage-specific promoters ^27^. We then sought to examined whether the candidates had potential to promote ZGA by activating MERVL and 2C-genes (Supplementary Fig. 1a and Supplementary Table 1-2). We constructed a 2C::tdTomato reporter mESC line with transgenic tdTomato red fluorescence protein driven by MERVL 5’-LTR. ^28^ (Fig. 2a). Candidate genes were cloned and overexpressed in the reporter cell line (Supplementary Fig. 1b-c). Interestingly, ectopic expression of *Obox4* markedly induced tdTomato expression, similar to that induced by *Dux* (Fig. 2b and Supplementary Fig. 2a-b). To characterize the impact of *Obox4* expression on 2C-genes, we established an ESC stable line bearing an inducible *Obox4* transgene (Fig. 2c). Differentially expressed gene analysis revealed that *Obox4* induction led to transcriptome changes in mESCs, characterized by upregulation of 2C-genes and TEs that were highly expressed in early-to-middle 2C-embryos ^29^ (Fig. 2d-e), naturally occurred 2C-like mESCs ^30^, and *Dux* induced 2C-like mESCs ^16^ (Fig. 2f, Supplementary Table 3). A substantial fraction (159/164) of *Obox4* induced 2C genes were also induced by *Dux* (Fig. 2g), while transcriptomic perturbance induced by *Obox4* is milder comparing with *Dux*, characterized by less number and less fold-upregulation of 2C genes (Fig. 2g-h). Interestingly, Dux and Obox4 were mutually inductive, where one promoted expression of the other (Fig. 2h). These results suggest that *Obox4* is an inducer of 2C-like genes and potentially redundant factor of *Dux*.

**Figure 2.**
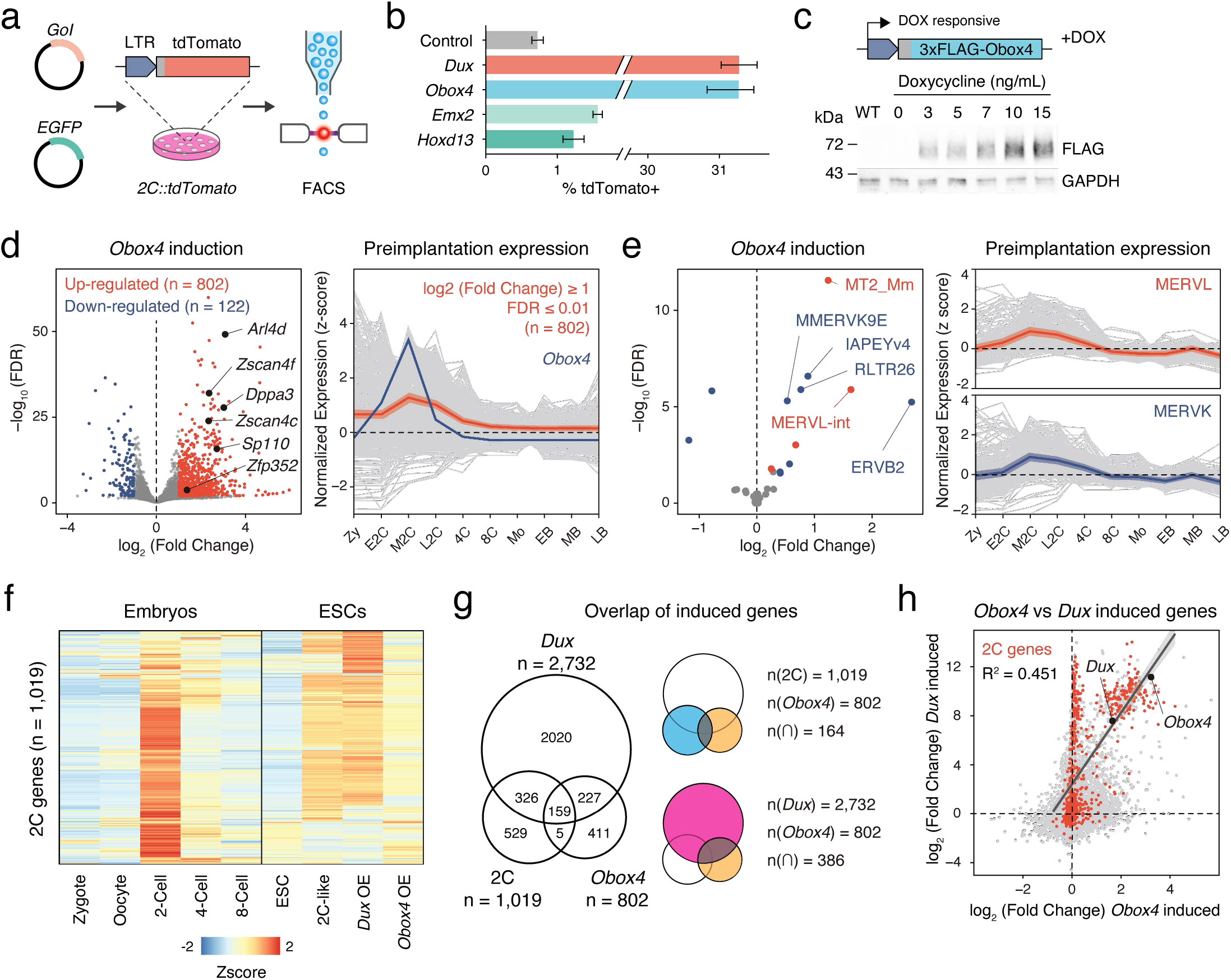
*Obox4* and *Dux* induce 2C-gene expression in mESCs. a) Diagram of the 2C::tdTomato reporter assay. ESCs bearing the tdTomato expression cassette under the control of the MERVL LTR promoter showed red fluorescence upon entering the 2C-like state. The expression of the transcription factor increased the 2C-like population, as detected using FACS. An EGFP expression plasmid was co-transfected with a gene of interest, to normalize the transfection efficiency. b) Boxplot showing normalized 2C-like cell percentage in 2C::tdTomato reporter ESCs overexpressing candidate pioneer factors. *Dux* and *Obox4* potently induced a 2C-like state. c) Upper panel: schematic of *Obox4*-inducible cell line construction. Lower panel: western blot showing OBOX4 level upon induction by different concentrations of doxycycline. Expression of OBOX4 was carried out in a dose-dependent manner. d) Left panel: volcano plot of DEGs in mESCs with *Obox4* induction for 48 hours. Representative 2C-genes are labeled with gene symbols. Right panel: expression profile of genes up-regulated by *Obox4* during embryogenesis. e) Left panel: volcano plot of differentially expressed transposable elements in mESCs with *Obox4* induction for 48 hours. MERVL and MERVK elements were highlighted. Right panel: expression profile of MERVL and MERVK elements during pre-implantation embryogenesis. f) Heatmaps of the expression of 2C-genes in preimplantation embryos, naturally occurred 2C-like mESCs, and induced 2C-like mESCs. g) Venn diagram showing overlap of 2C-genes with genes induced by ectopic expression of *Dux* and *Obox4* in mESCs. h) Scatterplot showing per-gene expression changes in *Dux* induced mESCs *versus Obox4* induced mESCs. 2C-genes are highlighted in red. *Obox4* and *Dux* are labeled.

### Obox4 binds to 2C-gene promoters and LTR elements in mESC

*Obox4* contains a homeobox, activates 2C-genes and TEs, and upregulates *Dux*-induced genes, which prompted us to examine whether OBOX4 is a transcription factor that activates 2C-gene associated loci through direct binding with *cis*-regulatory elements. Cleavage under targets and release using nuclease (CUT&RUN) leverages antibody-targeted cleavage of proximal DNA to identify the binding sites of DNA-associated proteins ^31^. To examine this technique, we performed CUT&RUN against triple FLAG-tagged DUX (3×FLAG-DUX) expressed in mESCs using a high-affinity anti-FLAG antibody ^32^, and confirmed that the DUX binding pattern revealed by CUT&RUN recapitulated the published HA-tagged DUX ChIP data ^16^ (Supplementary Fig. 3a-b). We then proceeded with characterizing genomic footprint of OBOX4 by performing CUT&RUN against 3×FLAG-OBOX4 expressed in mESCs, and discovered ∼24,000 peaks, among which 39.5% located in the gene promoter regions that covered 26.8% (273/1,019) of the 2C-genes (Fig. 3a). *De novo* motif discovery using the top 500 CUT&RUN signal peaks predicted CTGGGATYWRMR as top OBOX4 binding motif, which is enriched in the promoter regions of 2C-genes (Fig. 3b). Collectively, OBOX4 and DUX targeted 48.1% (490/1,019) of the 2C-genes with a considerable overlap (36.3% for OBOX4 and 31.3% for DUX), which covered many important ZGA genes including *Dppa2*, *Sp110*, and *Zscan4d* (Fig. 3c-d and Supplementary Table 4). The overlap was substantiated by the observation that MERVL LTR MT2_Mm was among the top 10 LTR targets of both OBOX4 and DUX, based on loci coverage (Fig. 3e-f and Supplementary Table 5). Notably, while DUX strongly prefers MT2_Mm, OBOX4 binding was biased toward MERVK LTRs, namely RLTR9 and RLTR13 elements (Fig. 3g), which also demonstrated a 2C-specific expression profile (Supplementary Fig. 3c-d). To examine whether these bindings were functional in the absence of *Dux*, we generated Obox4/Dux single and double knockout mECS lines, in which 14 of 15 protein-coding *Obox4* copies were removed from the genome (Supplementary Fig. 4-5). tdTomato reporters driven by MT2_Mm and RLTR13B2 were co-transfected with *Dux* and *Obox4* into *Obox4*/*Dux* double knockout ESCs (Fig. 3h). Analysis of the tdTomato-positive population showed that *Obox4* activated both RLTR13B2 and MT2_Mm, whereas *Dux* only activated MT2_Mm (Fig. 3i). These observations demonstrated that OBOX4 binds and activates a subset of DUX targets in mESCs and redundantly drives their expression in the absence of DUX.

**Figure 3.**
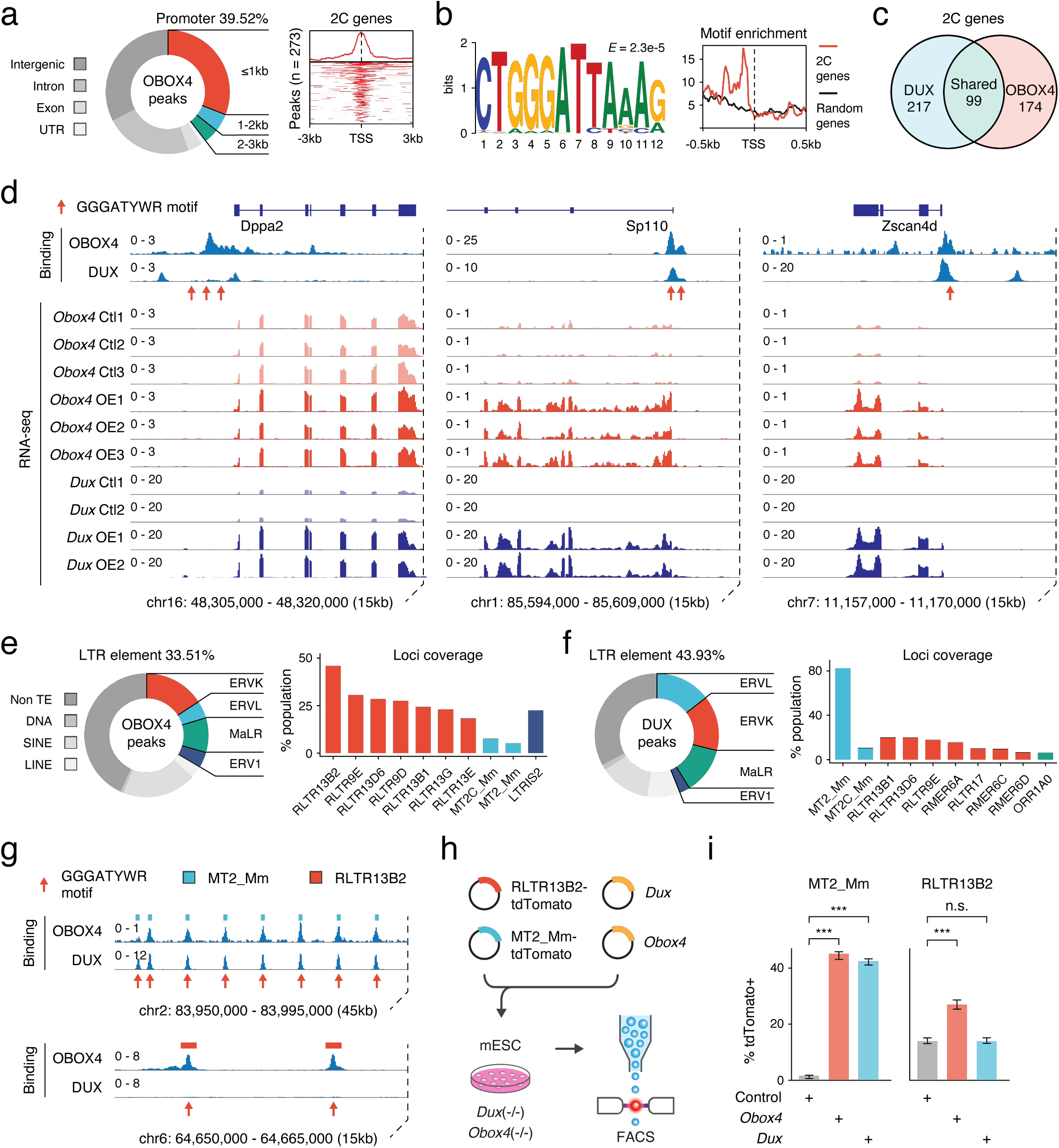
OBOX4 binds and activates 2C-specific LTR elements. a) Left panel: pie-chart displaying proportions of annotated genomic regions of the OBOX4 binding sites. Right panel: heatmap showing the OBOX4 binding site distribution near 2C-gene promoters. b) Left panel: the predicted OBOX4 binding motif using the top 500 CUT&RUN peaks. Right panel: histograph showing the distribution of predicted OBOX4 binding motif near 2C and random gene promoters. c) Venn diagram showing the distinct and overlap 2C-genes targeted by DUX and OBOX4. d) Representative genomic track showing DUX and OBOX4 binding sites at the *Dppa2*, *Sp110,* and *Zscan4d* loci and their expression levels in *Obox4* and *Dux* induced mESCs. Read counts were CPM normalized. The OBOX4 binding sites overlapped with those of DUX. *Dppa2, Sp110*, and *Zscan4d* expression was upregulated upon *Obox4* and *Dux* induction. e) Left panel: pie-chart displaying proportions of annotated TEs of the OBOX4 binding sites. Right panel: Bar plot showing the top 10 OBOX4 covered LTR elements. f) Left panel: pie-chart displaying proportions of annotated TEs of the DUX binding sites. Right panel: Bar plot showing the top 10 DUX covered LTR elements. g) Representative genomic track showing DUX and OBOX4 binding sites at MT2_Mm and RLTR13B2. Read counts were CPM normalized. The OBOX4 binding sites overlapped with those of DUX at MT2_Mm loci, while RLTR13B2 was uniquely bound by OBOX4. h) Schematic design of the LTR::tdTomato reporter assay. Plasmids bearing a tdTomato ORF downstream of MT2_Mm or RLTR13B2 were co-transfected with *Dux* or *Obox4* expression plasmids. Activation of LTR elements resulted in an increased red fluorescence-positive mESC population, which was measured using FACS. EGFP expression plasmid was co-transfected into the culture, following which the green fluorescence-positive population was measured, to normalize the transfection efficiency. i) Bar plots showing the percentage of red fluorescence-positive mESCs upon expression of *Dux* or *Obox4*. Both *Dux* and *Obox4* activated MT2_Mm, whereas only *Obox4* activated RLTR13B2.

### Concomitant loss of *Obox4* and *Dux* impairs preimplantation development

We then sought to determine whether *Obox4* is functionally required for pre-implantation development, particularly in the absence of *Dux*. Transient depletion of DUX has previously confused the field in that it caused an embryonic phototype while subsequent *Dux* knockout females were shown to be fertile ^15,18–21^. Therefore, it is critical to examine the functional requirement of OBOX4 and DUX in genetic knockout models. Somatic cell nuclear transfer (SCNT) is a technique to create embryos by transferring nuclei of somatic cells into enucleated oocytes (Fig. 4a), which recapitulates ZGA ^33^. When the knockout mESCs were subjected to SCNT as nuclear donors, 54.4% of WT mESC derived embryos developed to blastocyst stage at 4 days post nuclear transfer (dpt). *Obox4* and *Dux* single knockout donors led to reduced blastocyst formation rate at 38.5% and 39.7% respectively, whereas double knockout resulted in additive effect that led to significantly lowered blastocyst rate of 29.3% (Fig. 4b). Notably, all the blastocysts derived from double knockout mESC were of low quality, judging from severe morphological abnormality (Fig. 4c).

As an alternative genetic knockout model, we generated mice bearing *Dux* and *Obox4* knockout allele by direct CRISPR-Cas9 editing in embryos (Fig. 4d and Supplementary Fig. 4a-b). Among 39 *Obox4* copies, there are 15 copies that maintain intact open reading frame (ORF) for full-length OBOX4 (Supplementary Fig. 4c). While 14 of the 15 intact ORFs form tightly packed cluster (*Obox4* cluster), a solo ORF (*Obox4-ps33*) remains distant from the *Obox4* cluster and is interspersed with *Obox1*/*2*/*3*, a collection of *Obox* family members that are critical for ZGA ^34^. To minimize collateral genetic toxicity and interference to experiment result caused by removal of other *Obox* members, only *Obox4* cluster was knocked out and the solo ORF was retained (Supplementary Fig. 4d and Supplementary Fig. 6). The observed *Obox4*^KO^ frequency (7/29) in the progenies of *Obox4*^Het^ × *Obox4*^Het^ crossing is consistent with the expected 25% Mendelian ratio (Fig. 4e, Supplementary Fig. 7, and Supplementary Table 6). That *Obox4*^KO^ mice developed to adulthood without discernable abnormalities and are fertile when intercrossed suggest that development is compatible with loss of *Obox4* (Fig. 4f-g). We next attempted to produce *Dux*/*Obox4* double knockout (*Dux*^KO^/*Obox4*^KO^) mice to examine whether concomitant loss of *Dux* and *Obox4* compromises embryogenesis. We genotyped 54 pups from three litters of *Dux*^KO^/*Obox4*^Het^ × *Dux*^KO^/*Obox4*^Het^ and six litters of *Dux*^KO^/*Obox4*^Het^ × *Dux*^Het^/*Obox4*^Het^ mating pairs (Supplementary Fig. 8 and Supplementary Table 6). One *Obox4*^KO^/*Dux*^KO^ pup was present, and the frequency of *Obox4*^KO^/*Dux*^KO^ and *Obox4*^KO^/*Dux*^Het^ were significantly lower than the expected 25% Mendelian ratio (Fig. 4h). Development monitoring and genotyping of embryos produced by *Dux*^KO^/*Obox4*^Het^ × *Dux*^Het^/*Obox4*^Het^ mating pairs at 4.5 days post cotium (dpc) revealed that *Dux*^KO^/*Obox4*^KO^ was under-represented in blastocysts while over-represented in 2-cell arrest and degenerated embryos (Fig. 4i, Supplementary Fig. 9a-c, and Supplementary Table 7). Single blastomere genotyping and RNA-seq of *Dux*^KO^/*Obox4*^KO^ 2C embryos (Supplementary Fig. 9d-e) showed dysregulation of 2C-genes and TEs targeted by OBOX4 (Fig. 4j). The impaired development and transcriptome of *Dux*^KO^/*Obox4*^KO^ embryos at 2-cell stage showed that ZGA was defective in those embryos.

**Figure 4.**
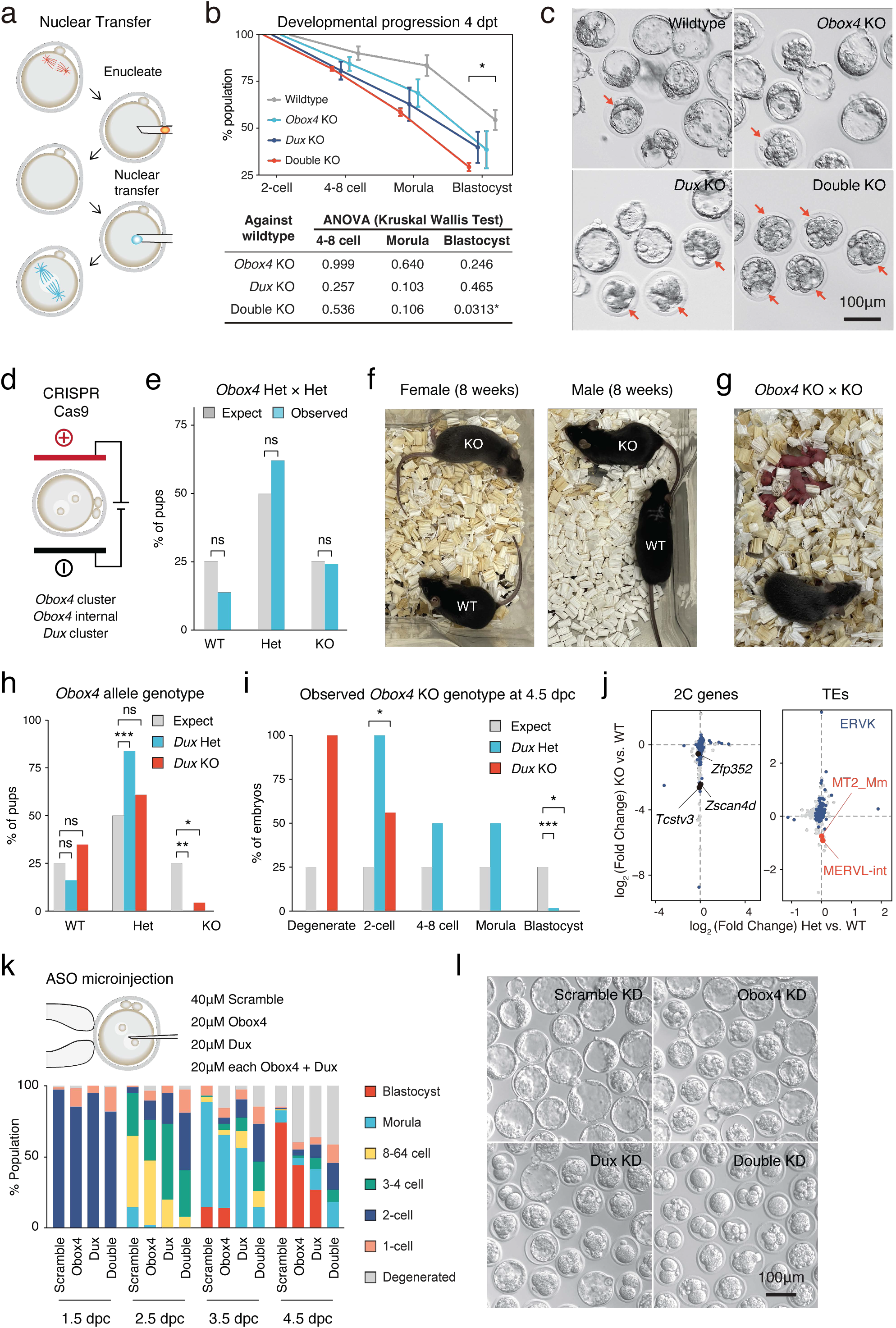
Concomitant loss of OBOX4 and DUX severely hinders ZGA. a) Schematic representation of somatic cell nuclear transfer (SCNT) experiment. The nuclei of knockout mESCs were transferred to enucleated oocytes to generate zygotes with knockout genotype. b) Upper panel: percent SCNT embryos developed to different stages at 4 dpt. Lower panel: *p*-value of non-parametric analysis of variance (ANOVA) among different genotypes and stages comparing to wildtype SCNT embryos. Three independent experiments were conducted, with 150-200 embryos per condition. c) Representative picture of SCNT embryos at 4 dpt. Morphologically abnormal blastocysts are highlighted. Blastocysts generated by double knockout mESCs were severely defective. d) Schematic representation of CRISPR-Cas9 mediated *Dux* and *Obox4* knockout mouse production. *In vitro* fertilized mouse zygotes were electroporated with pre-assembled CRISPR-Cas9 complex targeting *Dux* and *Obox4* loci. e) Bar plot showing genotype percentage of the pups delivered by *Obox4*^Het^ intercrosses. 4 litters delivered 29 pups, litter size 7.25 ± 1.26. ns *p*-value = 0.9146, chi-square goodness of fit test. f) Representative photos of *Obox4*^KO^ and WT adult mice analyzed in **e**. g) Photo of *Obox4*^KO^ intercross litter with live pups. h) Bar plot showing genotype of *Obox4* allele in the pups delivered by crossing of *Dux*^KO^/*Obox4*^Het^ × *Dux*^KO^/*Obox4*^Het^ or *Dux*^KO^/*Obox4*^Het^ × *Dux*^Het^/*Obox4*^Het^. Nine litters delivered 54 pups, *Dux* heterozygous and knockout allele were present in 31 and 23 pups, respectively. ** *p*-value = 0.001306; * *p*-value = 0.02218; chi-square goodness of fit test. i) Bar plot showing observed percentages of different preimplantation stage embryos bearing *Obox4* KO allele with *Dux* heterozygous or knockout allele at 4.5 dpc. Among the total 94 embryos assessed, two degenerated, ten 2-cell arrest, two 4-8 cell arrest, four morula arrest embryos were observed at 4.5 dpc, whereas 76 developed to blastocyst. *** *p*-value = 3.564×10^-5^; for 2-cell * *p*-value = 0.04348; for blastocyst * *p*-value = 0.02549; chi-square goodness of fit test. j) Scatterplot showing expression log2 fold changes of genes (left panel) and TEs (right panel) in *Dux*^KO^/*Obox4*^KO^ *versus Dux*^KO^/*Obox4*^Het^ 2C embryos. 2C-genes targeted by OBOX4 and ERVK elements are highlighted in blue. MT2_Mm and MERVL-int are labeled. k) Upper panel: schematic representation of *Dux* and *Obox4* knockdown experiments in pre-implantation embryos. Male pronuclei of zygotes were microinjected with ASO targeting *Dux* or *Obox4* transcripts. Lower panel: the percentages of embryonic stages observed at 1.5 dpc, 2.5 dpc, 3.5 dpc, and 4.5 dpc. l) Representative picture of KD embryos at 4.5 dpc. No blastocyst was observed among double ASO knockdown embryos.

As a transient depletion model in addition to genetic knockout approaches, microinjection of antisense oligonucleotide (ASO) into male pronuclei of zygotes was performed to knockdown *Obox4* and *Dux* (Fig. 4k and Supplementary Fig. 10a-c). Monitoring development until 4.5 dpc showed that *Obox4* single knockdown resulted in moderate developmental retardation, with almost 50% of embryos reaching the blastocyst stage. Similarly, blastocyst formation was preserved in nearly 25% of *Dux* single knockdown embryos, which is consistent with previously reported *Dux* knockdown/knockout experiments ^15,18–21^. The *Obox4*/*Dux* double knockdown markedly compromised blastocyst formation, resulted in more than 80% embryos degenerated or arrested before the 4C stage, and less than 20% manifested morula-like morphology (Fig. 4k-l). The double knockdown embryos with morula-like morphology manifest dysregulated transcriptome characterized by failure of expressing morula specific genes and activation of apoptosis related pathways, suggesting that the development was dysfunctional instead of delayed (Supplementary Fig. 10e-g).

The consistency among constitutive knockout in SCNT embryos and living mice and ASO-mediated transient depletion demonstrated that the expression of *Obox4* and *Dux* is collectively important for pre-implantation development, and that *Obox4* is capable of promoting ZGA in a *Dux*-independent manner. Collectively, these data show that *Obox4* promotes mouse preimplantation development in the absence of *Dux*.

### OBOX4 promotes 2C-gene expression upon depletion of DUX

As *Obox4* has been shown to activate 2C-genes and TEs in mESCs, and embryonic depletion of OBOX4 impairs 2C-genes and TEs activation, we questioned whether the impairment can be ameliorated by restoration of OBOX4. First, we determined whether the presence of OBOX4 can rescue the developmental arrest of *Obox4*/*Dux* double knockdown embryos. Codon-optimized *Obox4* mRNA without the ASO target motif was produced using *in vitro* transcription. The rescue experiment was performed by means of co-microinjection of *Obox4* mRNA with *Obox4*/*Dux* double knockdown ASOs (Fig. 5a). Restoration of OBOX4 in *Obox4* mRNA-microinjected embryos was confirmed using immunofluorescence staining at the 2C stage (Fig. 5b). As expected, development monitoring showed that *Obox4*/*Dux* double knockdown embryos failed blastocyst formation at 4.5 dpc, whereas among *Obox4* mRNA-rescued embryos, blastocyst formation was retained at a similar level to that observed in the *Dux* single knockdown experiment (Fig. 5c-d and Fig. 4k-l). RNA-seq of 2C-embryos revealed exacerbated dysregulation of 2C-genes and TEs by *Obox4*/*Dux* double knockdown comparing to single knockdown, and the dysregulated transcriptome of double knockdown was rescued by resupplying OBOX4, revealed by number of up and down-regulated genes (Fig. 5e-f). Differential expression analysis revealed that 58% (102/175) of the down-regulated 2C-genes in double knockdown embryos were rescued, including 2C stage markers and important ZGA factors (Fig. 5g-h). These results showed that that OBOX4 redundantly activates 2C gene to promote ZGA under DUX depletion.

**Figure 5.**
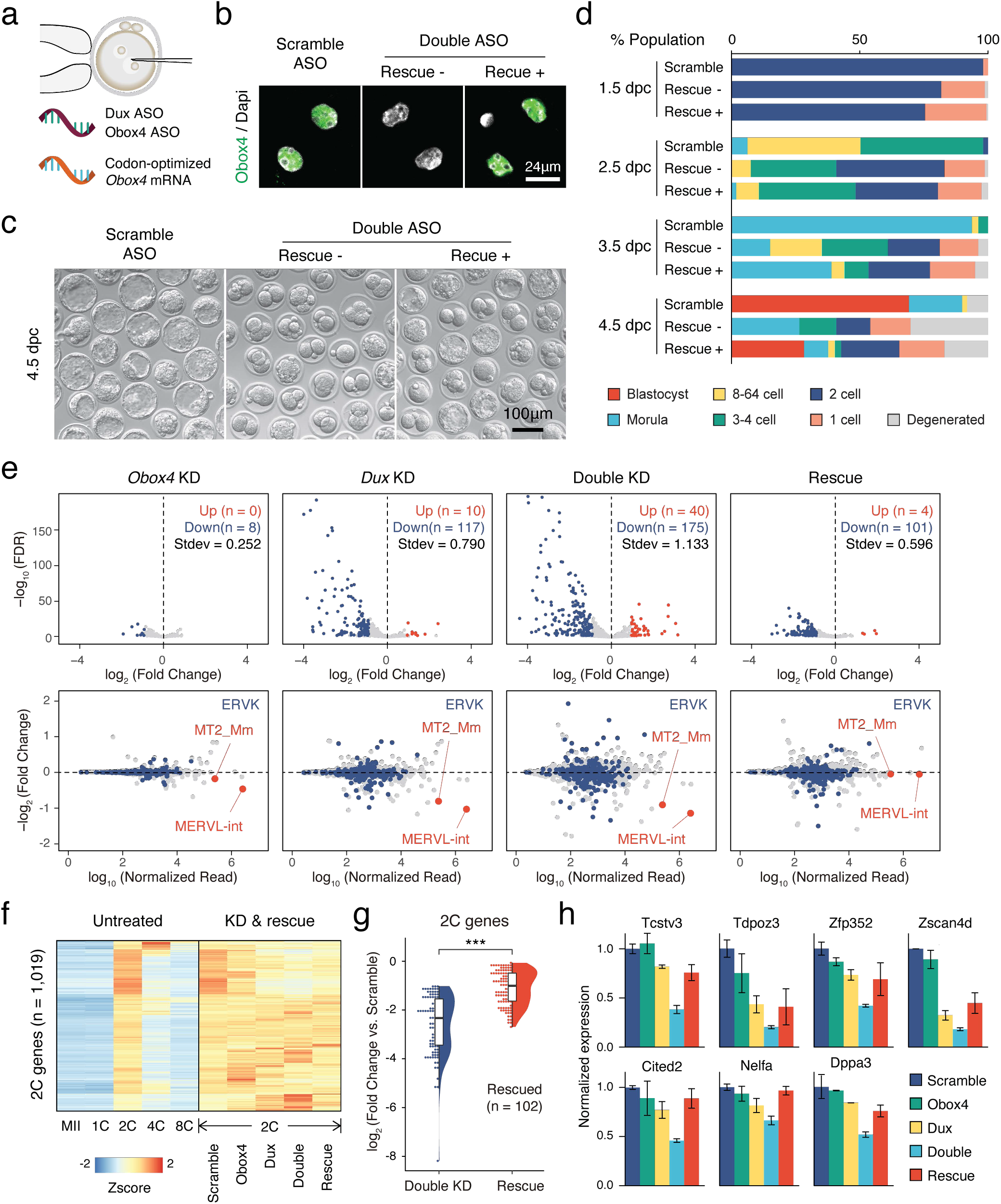
OBOX4 promotes ZGA in the absence of DUX. a) Schematic of the double knockdown rescue experiment. Male pronuclei of zygotes were injected with ASO targeting *Obox4* and *Dux* transcripts as well as *in vitro*-transcribed codon-optimized *Obox4* mRNA. b) Immunofluorescence staining of OBOX4 in early 2-cell embryos microinjected with scrambled ASO, double ASO (*Obox4* and *Dux*), or double ASO with codon-optimized *Obox4* mRNA. c) Representative picture of knockdown and rescue embryos at 4.5 dpc. Codon-optimized *Obox4* mRNA injection rescued blastocyst formation in ASO knockdown embryos. d) The percentages of embryonic stages observed at 1.5 dpc, 2.5 dpc, 3.5 dpc, and 4.5 dpc. The plot represents the sum of three independent experiments, with 80-100 embryos per condition. e) Upper panel: volcano plot showing the results of DEG analysis of 2C-genes in knockdown and rescue embryos, as compared to that in scramble ASO-injected embryos. Standard deviations of log2(fold-change) were used to represent the degree of transcriptome dysregulation. Stdev, standard deviation. Lower panel: MA plot of differential expression of TEs in knockdown and recue embryos, as compared to that in scramble ASO-injected embryos. MERVK elements and MERVL are highlighted. f) Heatmaps of the expression of 2C-genes in preimplantation embryos, knockdown 2C-embryos, and rescue 2C-embryos. g) Rain plot displaying the expression change distribution of rescued 2C-genes in double knockdown and recue 2C-embryos. h) Bar plots showing the expression of representative 2C-genes in knockdown and recue embryos; n=3 biological replicates.

## Discussion

Starting with a transcriptionally inert genome, the eutherian ZGA engages in massive yet coordinated expression of genes driven by LTR TEs. In mice, it has been reported that the transcription factor DUX accesses and opens condensed chromatin of the MERVL LTR loci, which results in the activation of downstream ZGA genes ^15–17^. However, the establishment of fertile *Dux* knockout mice suggests that ZGA is redundantly driven by other transcription factor(s) ^18–21^. In the present study, we found that *Obox4*, a cryptic multi-copy cluster gene, encodes a homeodomain-containing protein that potently induces ZGA gene expression. Using multiple genetic knockout models, we showed that concomitant depletion of *Dux* and *Obox4* was hardly compatible with embryogenesis, characterized by *Dux*/*Obox4* double knockout mESCs compromised SCNT and mouse produced at sub-Mendelian ratio. Consistently, *Obox4*/*Dux* double knockdown markedly compromised pre-implantation development, which tolerated single knockdown of either gene. Double knockdown embryos exhibited severely dysregulated transcriptomes, characterized by MERVL and ZGA gene activation failure. We also characterized the molecular mechanisms underlying the biological significance of *Obox4* as well as its relevance to *Dux*. OBOX4 directly binds to MERVL (the target of DUX) and MERVK loci and activates specific MERVL and MERVK elements in a *Dux*-independent manner. In summary, our results highlight that OBOX4 is a transcription factor that is functionally redundant to DUX during ZGA.

*Obox* transcripts were first discovered in gonads ^35^, and were later found to be highly abundant in mouse 2C-embryos ^36^. The *Obox* family has 67 members clustered in the sub-telomeric region of mouse chromosome 7 and is further divided into six subfamilies: *Obox1*, *Obox2*, *Obox3*, *Obox4*, *Obox5*, and *Obox6*, with *Obox4* constituting nearly 60% (39 of 67) of the family population ^37,38^. Although *Obox4* dominates the *Obox* family population, members of the entire subfamily have been annotated as pseudogenes and have largely been neglected from investigation. Despite being abundantly transcribed in the gonads, the biological significance of *Obox* genes is poorly understood. The role of *Obox4* is particularly mysterious, as it remains to be determined whether *Obox4* is a functional protein-coding gene. Transcription of *Obox4* was originally reported to be testis-specific and contradicted by the detection of *Obox4* transcripts in the ovaries and oocytes ^39^. *Obox4* is involved in oocyte maturation and ESC differentiation ^40–44^. However, these observations are supported by limited evidence, with no evidence at the protein level. By detecting OBOX4 in various totipotent cell types using high-quality monoclonal antibodies, we provided decisive evidence that *Obox4* is a functional protein-coding gene. Our data demonstrated that OBOX4 binds to genomic loci of MERVL-derived promoters and MERVK-derived enhancers and mediates their activation during ZGA. However, whether and how other *Obox* family members contribute to ZGA remains unclear.

It is worth noting that the ability of OBOX4 to promote ZGA appears to be weaker than that of DUX, characterized by lower fold upregulation of 2C-genes upon overexpression of OBOX4 than DUX, non-induction of MERVL Gag protein in mESCs, less dysregulated transcriptome in *Obox4* depleted embryos, and less adversarial phenotype in *Obox4* depleted embryos and mice. On the other hand, the severe cytotoxicity borne by DUX^45^ cannot be observed in OBOX4, as ectopic expression of OBOX4 in mESCs and embryos did not show hindrance in cell viability. Whether the variance in potency to promote ZGA has additional biological significance other than functional redundancy warrants further investigation.

Unlike *Dux*, which is structurally and functionally conserved throughout placentalia ^46^, the ancestral locus of the *Obox* family appears to have undergone mouse-specific duplication and generated a gene cluster that is collectively syntenic to the *Tprx2* locus in other mammals ^47^. Despite the distal homology between *Obox4* and *Tprx2*, it has been recently reported that human *TPRX2* is expressed in 8-cell embryos ^14^. Interestingly, TPRX2 was shown to cause defective ZGA upon embryonic depletion and bind important ZGA genes in hESCs ^48^, suggesting that the *Tprx2* locus has undergone functionally convergent evolution despite its divergent genetic context. However, whether *TPRX2* plays a similar role in humans to *Obox4* in mice regarding the redundancy to *Dux* (*DUX4* in humans) remains to be elucidated.

Genetic redundancy is widespread in higher organisms and typically arises in signaling networks, in which multiple functionally overlapping factors commit to a shared teleological objective, to counteract sporadic mutations or defects ^49,50^. This principle is of particular importance to embryogenesis, a process that gives rise to a reproducing organism. Indeed, genetic redundancy in pluripotency has been evidenced by the observation that naïve pluripotency can be elicited and maintained through various “independent inputs that operate through both unique and convergent targets”^51^. Although the molecular pathway that governs totipotency has been largely restricted to the scope that centers on *Dux*, it is reasonable to assume that the acquisition of totipotency also complies with the rule of redundancy. This assumption was validated by the generation of fertile *Dux* knockout mice; however, the panorama of ZGA redundancy remained hidden. Our work on *Obox4* sheds light on this topic by discovering that *Obox4* serves as a redundant gene to *Dux* during ZGA and is a bivalent activator of MERVL and MERVK elements. Yet that concomitant loss of *Dux* and *Obox4* is viable reveals that ZGA is so strongly canalized that many differences in genes make very little difference to the phenotype. Although the function of MERVL has been extensively investigated, particularly during early embryogenesis, the role of MERVK remains unclear. Interestingly, recent studies have suggested that MERVK serves as a meiosis-specific enhancer of spermatogenesis ^52^. Mammalian Y chromosomes have undergone rapid evolution and have been intensively inserted by TEs ^53^. As *Obox4* is highly expressed in the testes, it is worth investigating whether *Obox4* plays a regulatory role during spermatogenesis, by activating MERVK-derived enhancers.

Pioneer factors possess the ability to specifically recognize and access DNA motifs that are inaccessible to other transcription factors ^54^. In the scope of pluripotency-to-totipotency transition, OBOX4 and DUX are qualified pioneer factors for they are capable of accessing and activating silenced 2C-genes in ESCs. However, it is not likely that such pioneer activity demonstrated *in vitro* is accounted for ZGA initiation *in vivo*, considering that genome-wide demethylation precedes the timing of OBOX4 and DUX translation ^55^, and neither individual nor collective depletion of OBOX4 and DUX caused strict 2C arrest. Recent studies have identified the maternally deposited orphan nuclear receptor NR5A2 that triggers ZGA by activating short interspersed nuclear element (SINE) B1 family ^56,57^, while DNA damage induced p53 activation is important but dispensable for *Dux* activation in mouse zygotes ^58^. These results suggest another layer of genetic redundancy at ZGA initiation level. Hence, how *Dux* and *Obox4* are activated after fertilization and whether they share the same activation mechanism requires further interrogation.

## Supplemental figure legends

**Supplementary Figure 1.**
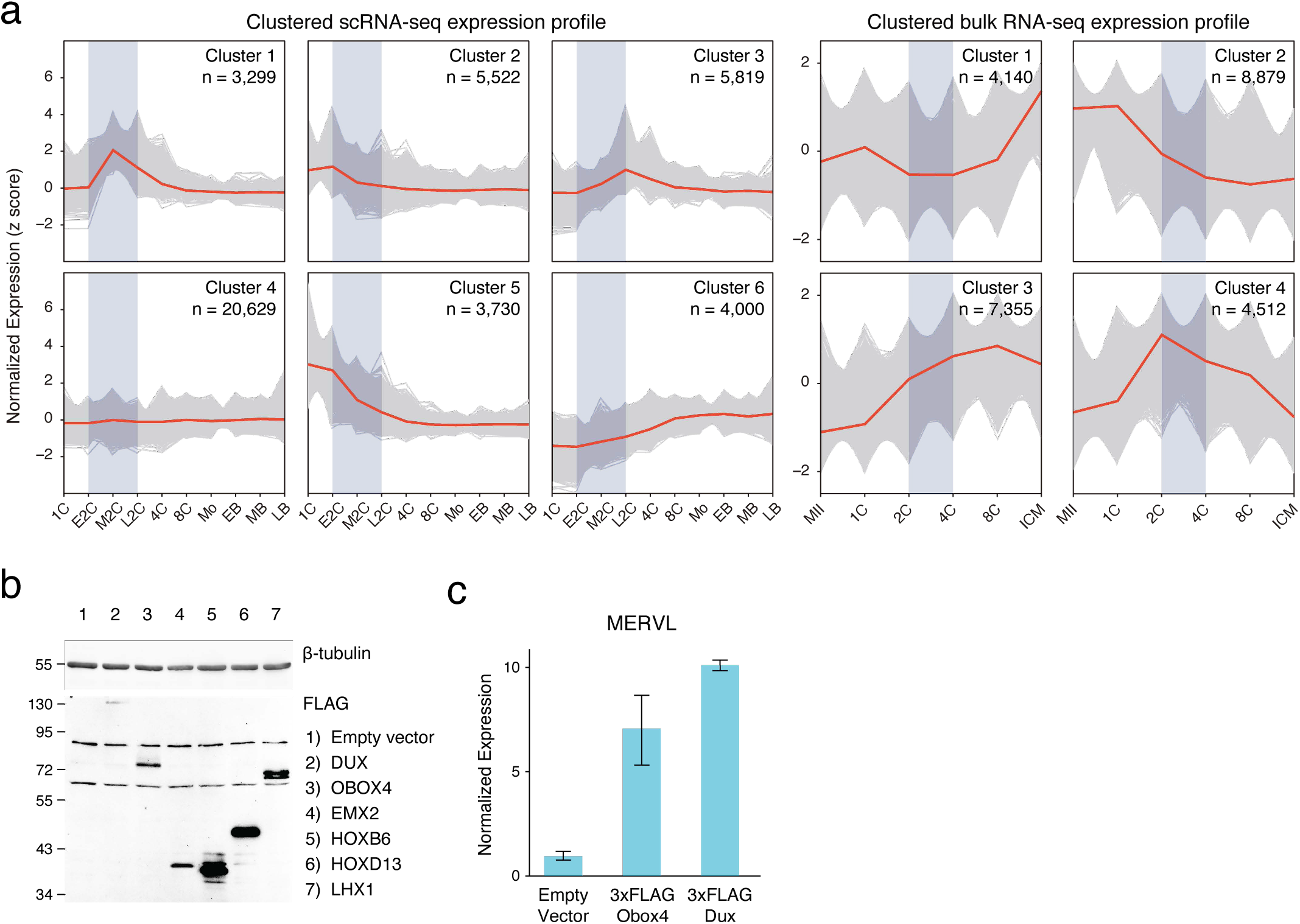
a) Expression profiles of *k*-means clustered gene sets during preimplantation development based on *z*-scored expression of scRNA-seq and bulk RNA-seq. Genes appeared in both scRNA-seq cluster 1 and bulk RNA-seq cluster 4 were designated as 2C-genes. b) Western blot showing protein products of the candidate genes ectopically expressed in mESCs at 18 hours post transfection. c) Bar plot showing MERVL transcript level in mESCs at 18 hours post *Dux* or *Obox4* transfection, detected by reverse transcription quantitative real-time PCR (RT-qPCR).

**Supplementary Figure 2.**
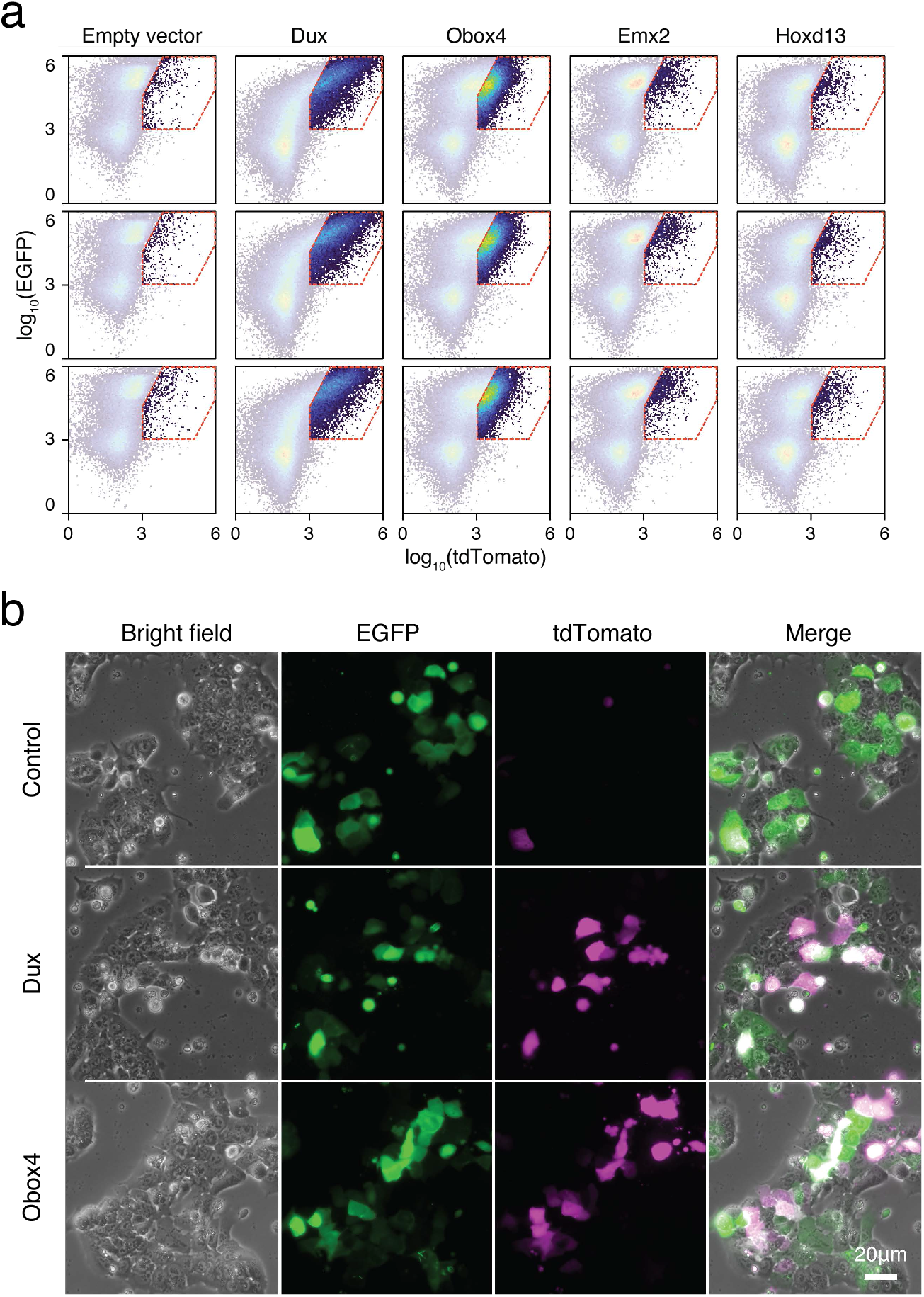
a) FACS analysis of 2C::tdTomato cells co-transfected with candidate gene expression plasmids and EGFP. b) Fluorescence microscopy of 2C::tdTomato ESCs transfected with empty plasmid, plasmid encoding DUX, or plasmid encoding OBOX4. Plasmids encoding EGFP were co-transfected to normalize the transfection efficiency in FACS analysis.

**Supplementary Figure 3.**
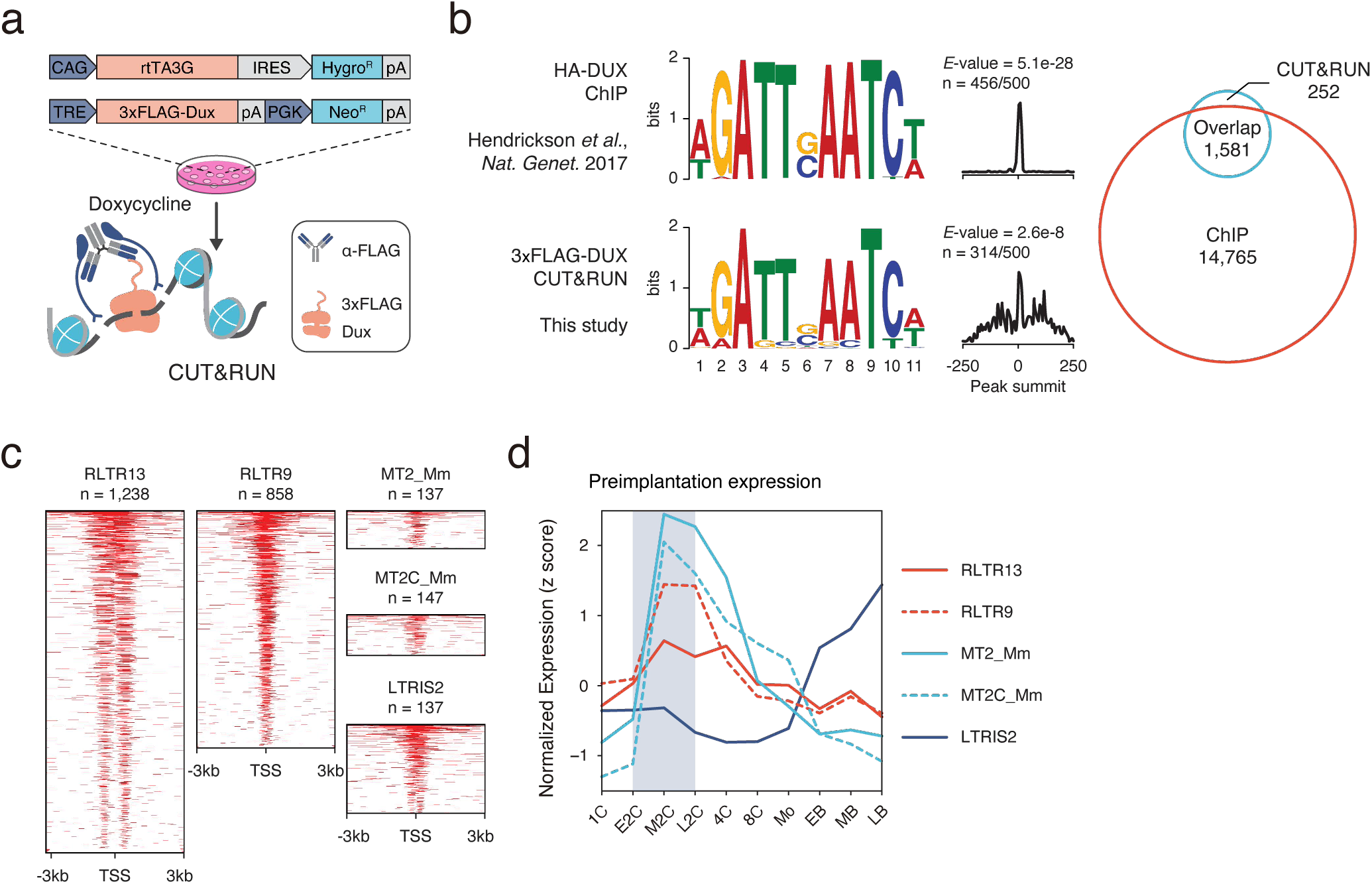
a) Schematic diagram of the CUT&RUN workflow using mESCs with induced expression of FLAG-tagged DUX. mESCs bearing a doxycycline-inducible 3×FLAG-OBOX4 expression cassette were induced with doxycycline for 24 h. OBOX4-associated DNA was pulled down in the CUT&RUN assay, using a high-affinity anti-FLAG antibody. b) Left panel: comparison of DUX binding motif predicted with published ChIP data and CUT&RUN result from this study. Right panel: overlap of DUX binding peaks discovered with the two data sets. c) Heatmaps showing the OBOX4 binding site distribution near top covered LTR elements. d) Expression profile of top 5 OBOX4 covered LTR elements during embryogenesis.

**Supplementary Figure 4.**
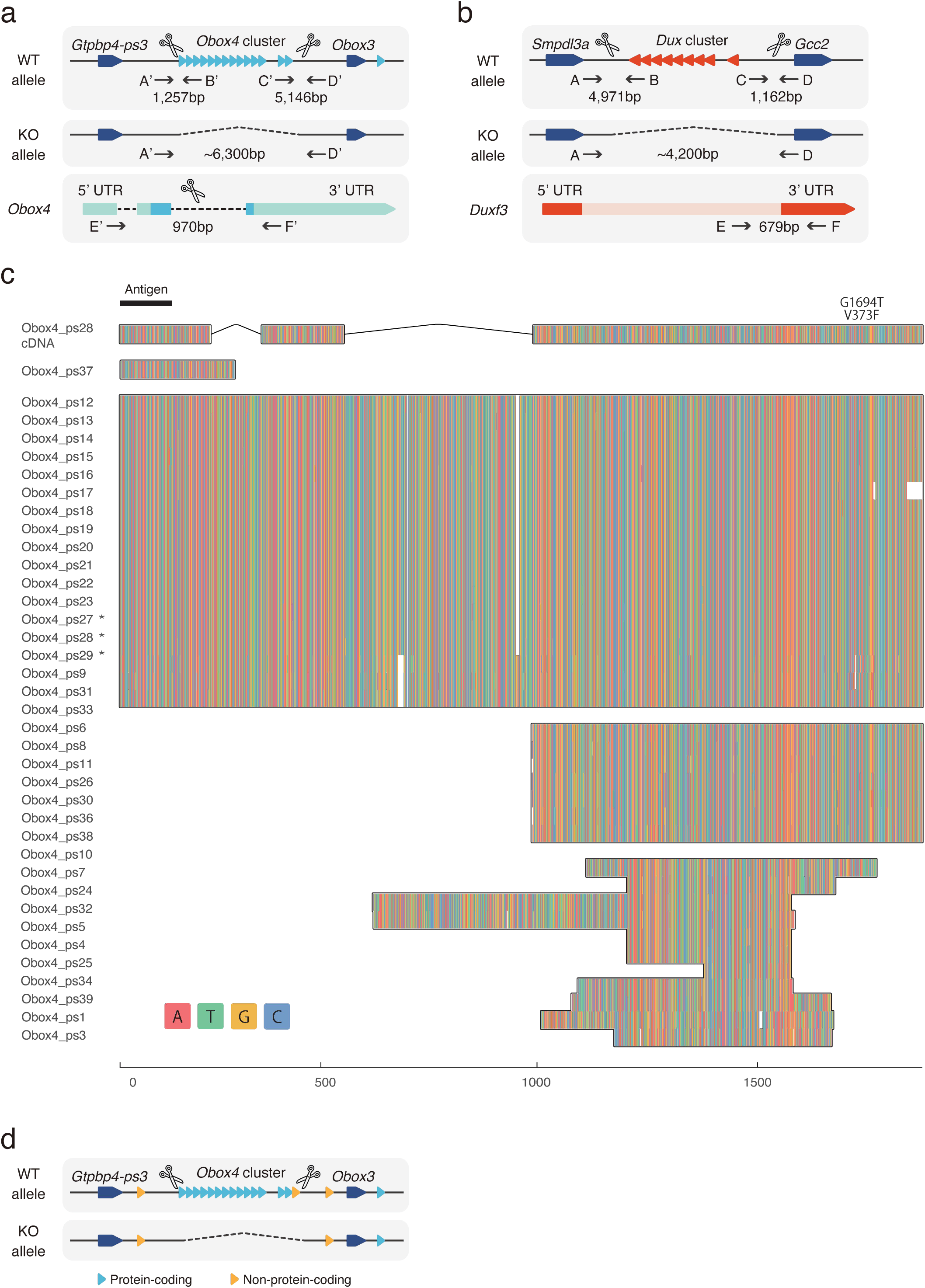
a) Schematic diagram of producing *Obox4* knockout mice using CRISPR-Cas9 (not drawn to scale). sgRNAs flanking the *Obox4* cluster as well as the internal sequence of individual *Obox4* were introduced into mouse zygotes with SpCas9, to knockout *Obox4*. Primers flanking Cas9 cutting sites (A’-D’) were designed to validate removal of target allele. Primers inside of *Obox4* (E’-F’) were designed to quantify *Obox4* copy number by qPCR. b) Schematic diagram of producing *Dux* knockout mice using CRISPR-Cas9 (not drawn to scale). Previously reported sgRNAs flanking the *Dux* cluster were introduced into mouse zygotes with SpCas9, to knockout *Dux*. Primers flanking Cas9 cutting sites (A-D) were used to validate removal of target allele. Primers inside of *Dux* (E-F) were designed to detect translocation of *Dux* cluster. c) Multiple sequence alignment of DNA sequence among the *Obox4* copies. Full length *Obox4* with internal top codons are marked by asterisks. d) Schematic diagram of *Obox4* cluster removal strategy and copy number quantification. Primer E’ and F’ targets all 15 protein-coding as well as 3 non-protein-coding *Obox4* loci, among which 14 protein-coding and 1 non-protein-coding copies were subjected to knockout.

**Supplementary Figure 5.**
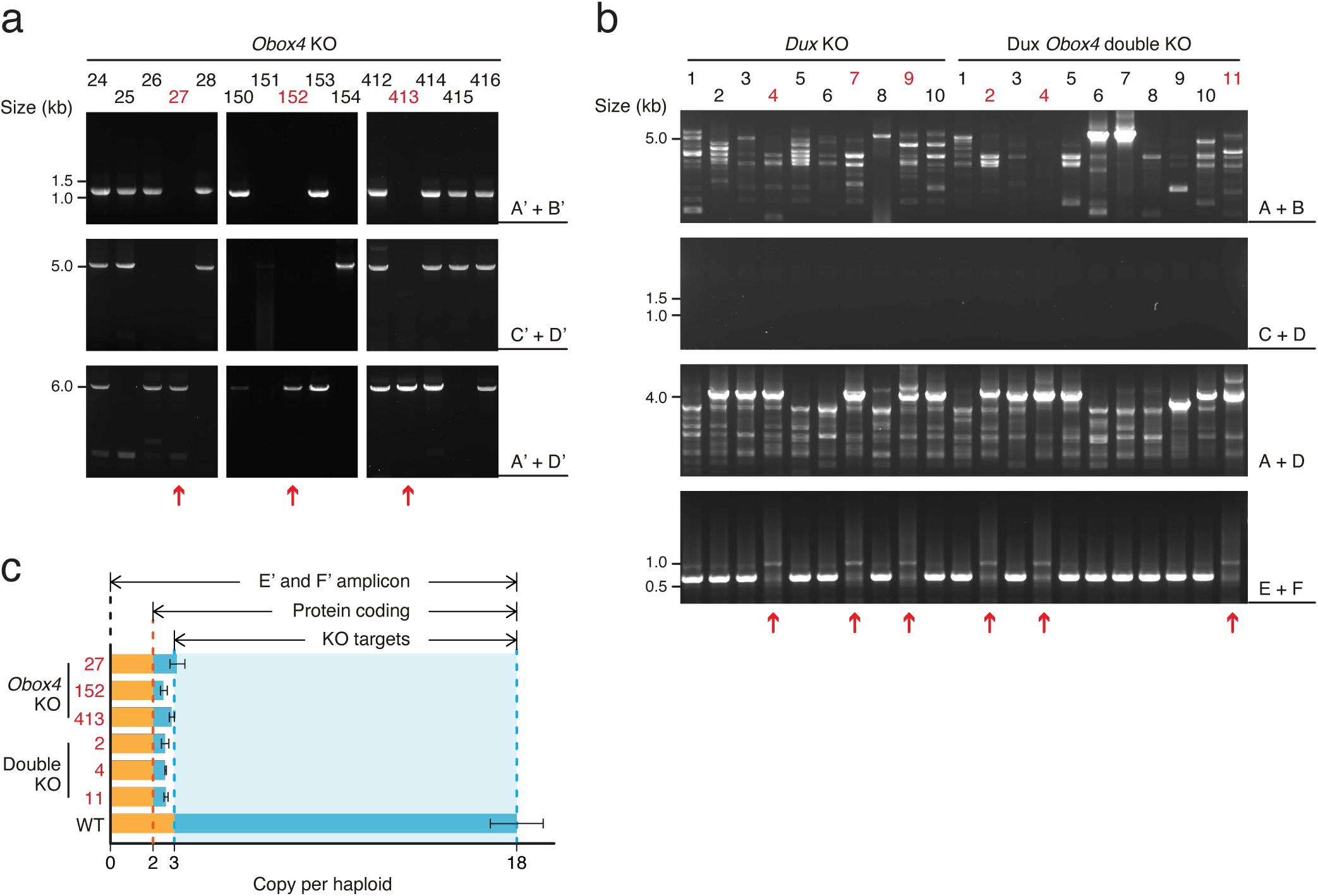
a) Agarose gel image of PCR-based genotyping of the *Obox4* knockout allele in founder mESC lines. The red arrows indicate cell lines bearing bi-allelic deletions of the *Obox4* cluster. b) Agarose gel image of PCR-based genotyping of the *Dux* knockout allele in founder mESC lines. The red arrows indicate cell lines without the genomic *Dux* sequences. c) Bar plot showing copy number of *Obox4* in the founder mESC lines analyzed in **a**-**b**, as detected by qPCR using purified genomic DNA.

**Supplementary Figure 6.**
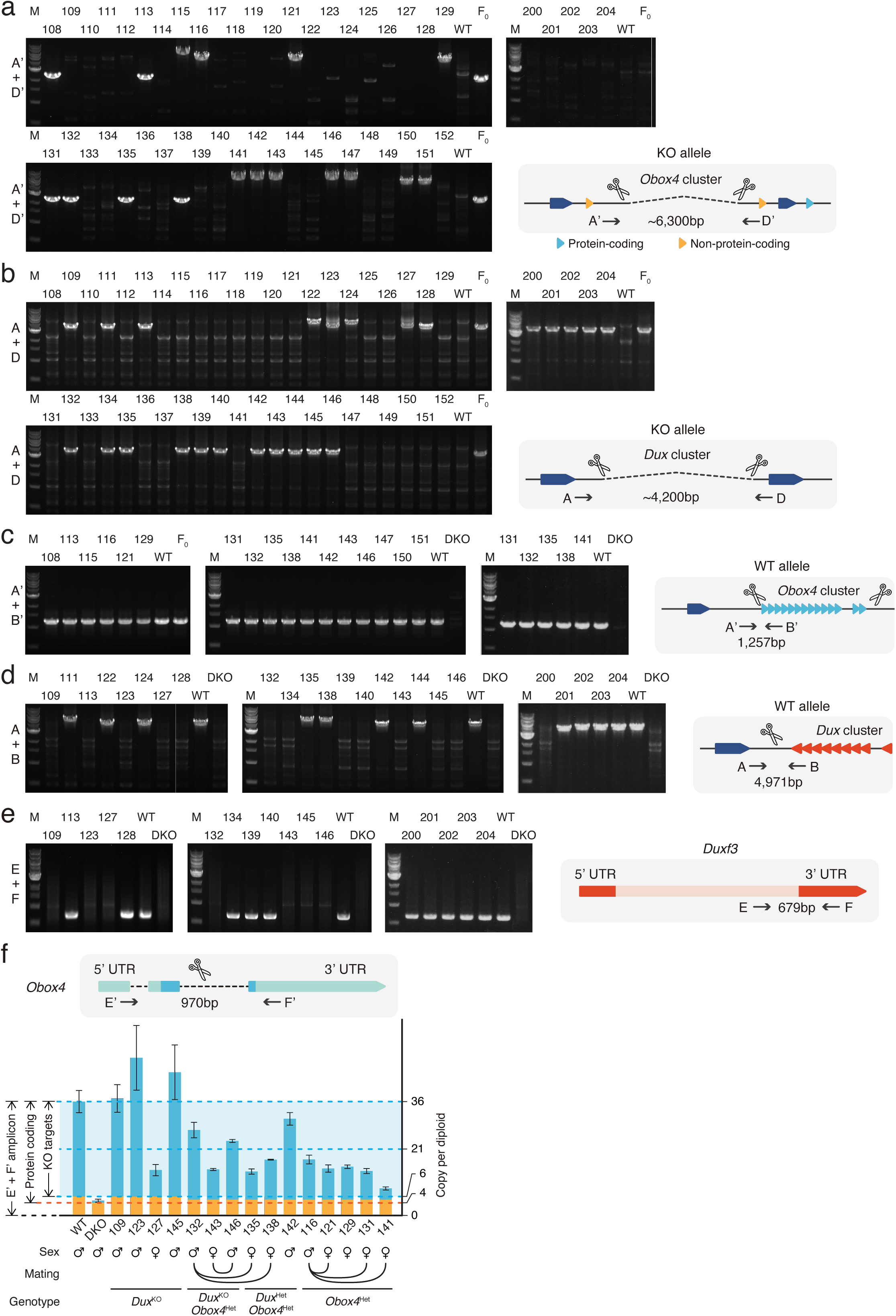
a) Agarose gel image of PCR-based genotyping of the *Obox4* knockout allele in the F1 mice. b) Agarose gel image of PCR-based genotyping of the *Dux* knockout allele in the F1 mice. c) Agarose gel image of PCR-based genotyping of the *Obox4* cluster 5’ cutting site allele in the F1 mice. d) Agarose gel image of PCR-based genotyping of the *Dux* cluster 5’ cutting site allele in the F1 mice. e) Agarose gel image of PCR-based genotyping of the *Dux* internal sequence in the F1 mice. f) Bar plot showing copy number of *Obox4* in the F1 mice analyzed in **a**-**e**, as detected by qPCR using purified genomic DNA.

**Supplementary Figure 7.**
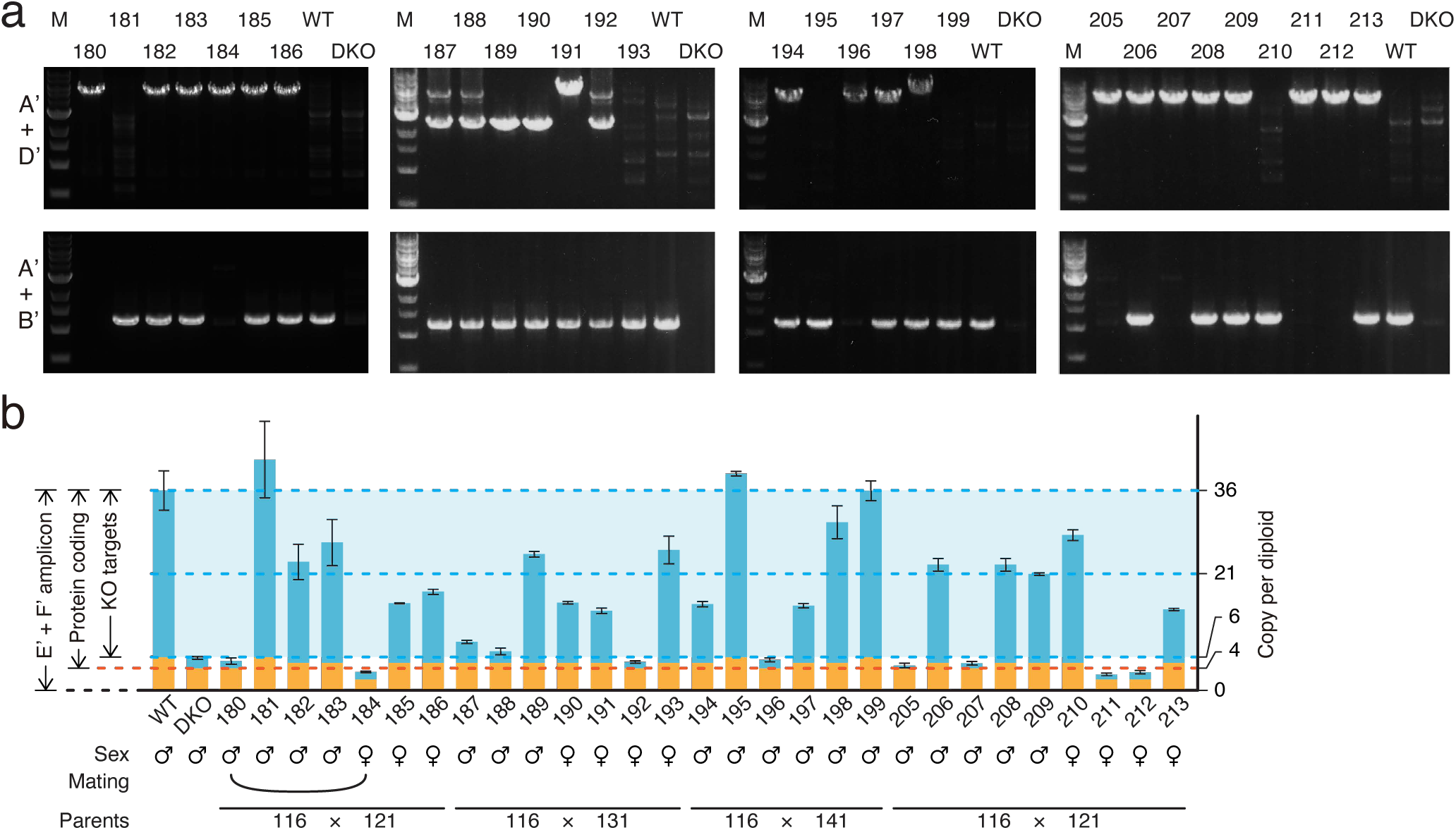
a) Agarose gel image of PCR-based genotyping of the *Obox4* knockout and *Obox4* cluster 5’ cutting site wild type allele in the F2 mice produced by intercrossing *Obox4*^Het^ F1 mice. b) Bar plot showing copy number of *Obox4* in the F2 mice analyzed in **a**, as detected by qPCR using purified genomic DNA.

**Supplementary Figure 8.**
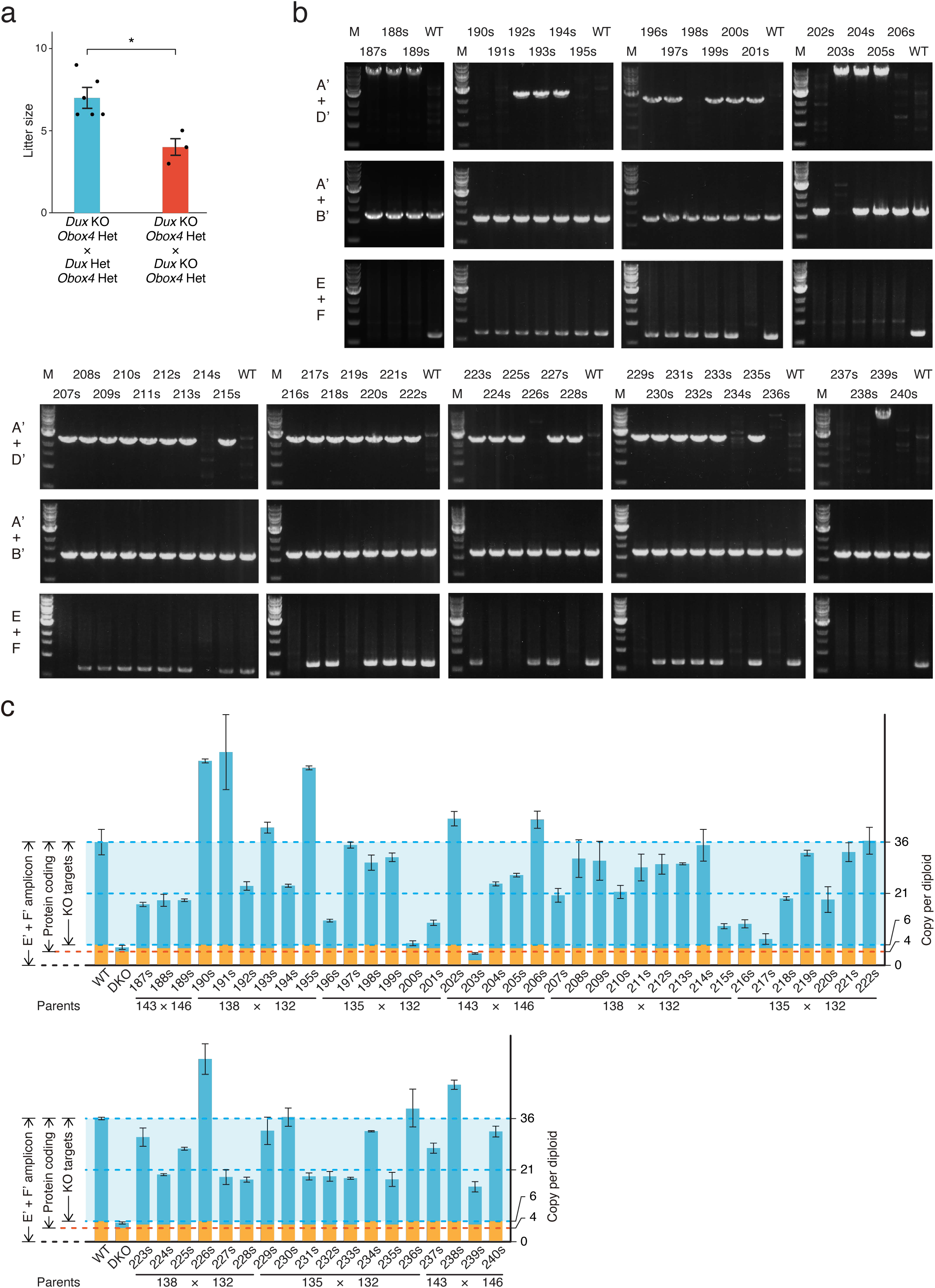
a) Bar and dot plot showing litter sizes of intercross used to produce *Dux*/*Obox4* double knockout mice. Each dot represents individual litter size. six and three litters were analyzed for *Dux*^KO^/*Obox4*^Het^ × *Dux*^Het^/*Obox4*^Het^ and *Dux*^KO^/*Obox4*^Het^ × *Dux*^KO^/*Obox4*^Het^ mating, respectively. Crossing of *Dux*^KO^/*Obox4*^Het^ × *Dux*^KO^/*Obox4*^Het^ delivered reduced litter size than those of *Dux*^KO^/*Obox4*^Het^ × *Dux*^Het^/*Obox4*^Het^. * *p*-value = 0.01104, two-tailed Student’s *t* test. Error bars indicate standard deviations. b) Agarose gel image of PCR-based genotyping of the *Obox4* knockout, *Obox4* cluster 5’ cutting site allele, and *Dux* internal sequence in the F2 mice produced by intercrossing *Dux*^KO^/*Obox4*^Het^ and/or *Dux*^Het^/*Obox4*^Het^ F1 mice. c) Bar plot showing copy number of *Obox4* in the F2 mice analyzed in **a**, as detected by qPCR using purified genomic DNA.

**Supplementary Figure 9.**
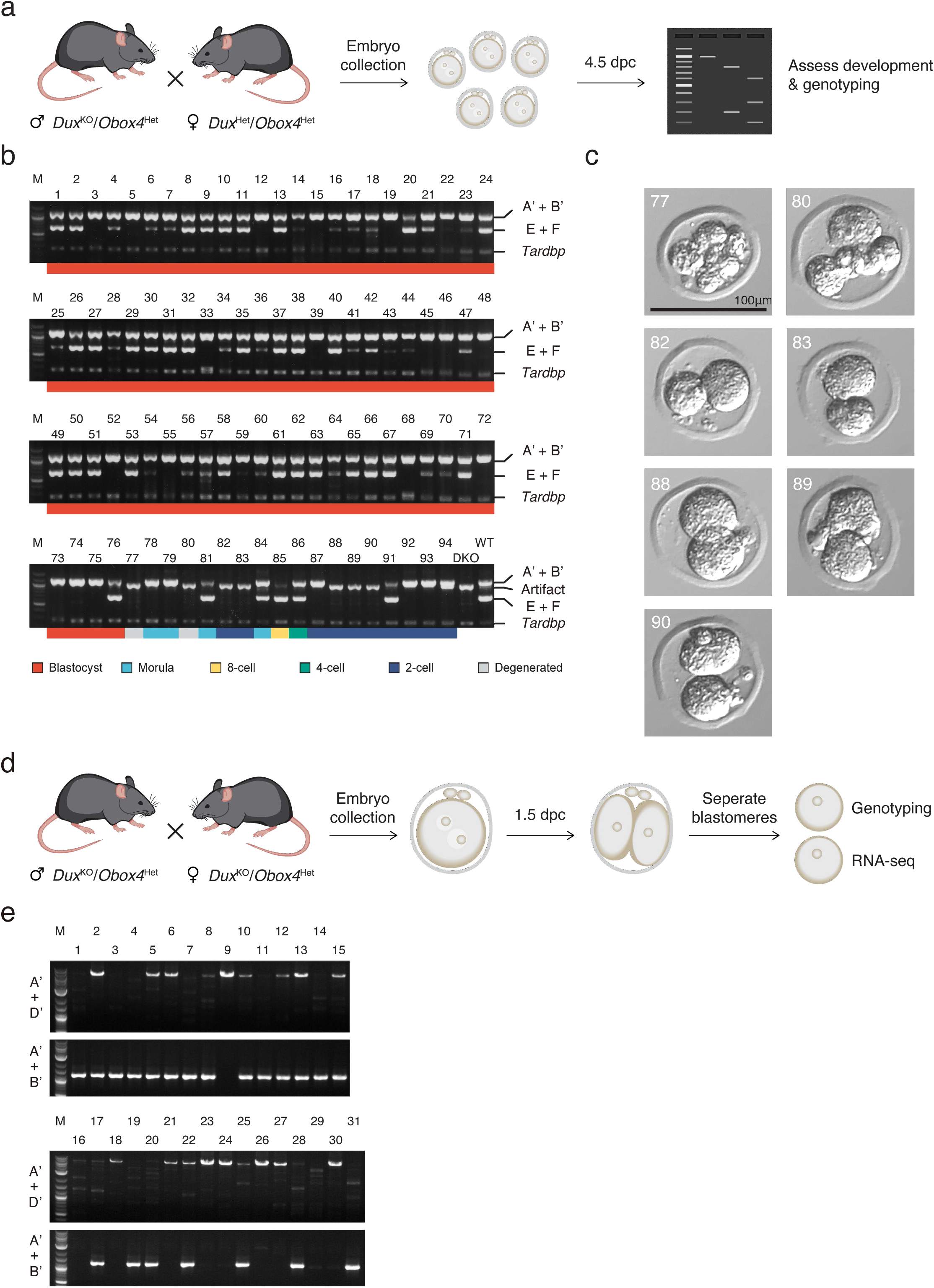
a) Schematic diagram of knockout embryo production for development monitoring. *Dux*^Het^/*Obox4*^Het^ female mice were mated with *Dux*^KO^/*Obox4*^Het^ male mice. Embryos were collected at 0.5 dpc and cultured *in vitro* until 4.5 dpc, followed by developmental assessment and genotyping. b) Agarose gel image of multiplex PCR-based genotyping of *Obox4* cluster 5’ cutting site allele, *Dux* internal sequence, and *Tardbp* internal sequence in the embryos produced by intercrossing *Dux*^KO^/*Obox4*^Het^ and *Dux*^Het^/*Obox4*^Het^ F1 mice. Note that multiplexing three sets of PCR primers produced artifact band in *Dux*^KO^/*Obox4*^KO^ embryos. c) Representative pictures of *Dux*^KO^/*Obox4*^KO^ embryos. d) Schematic diagram of knockout embryo production for RNA-seq. Embryos were collected at 0.5 dpc from *Dux*^KO^/*Obox4*^Het^ mating pairs and cultured *in vitro* until 2-cell stage at 1.5 dpc, blastomeres of each embryo were separated and subject to genotyping and RNA-seq respectively. e) Agarose gel image of genotyping of *Obox4* cluster 5’ cutting site allele in the embryos produced by intercrossing *Dux*^KO^/*Obox4*^He^ mice.

**Supplementary Figure 10.**
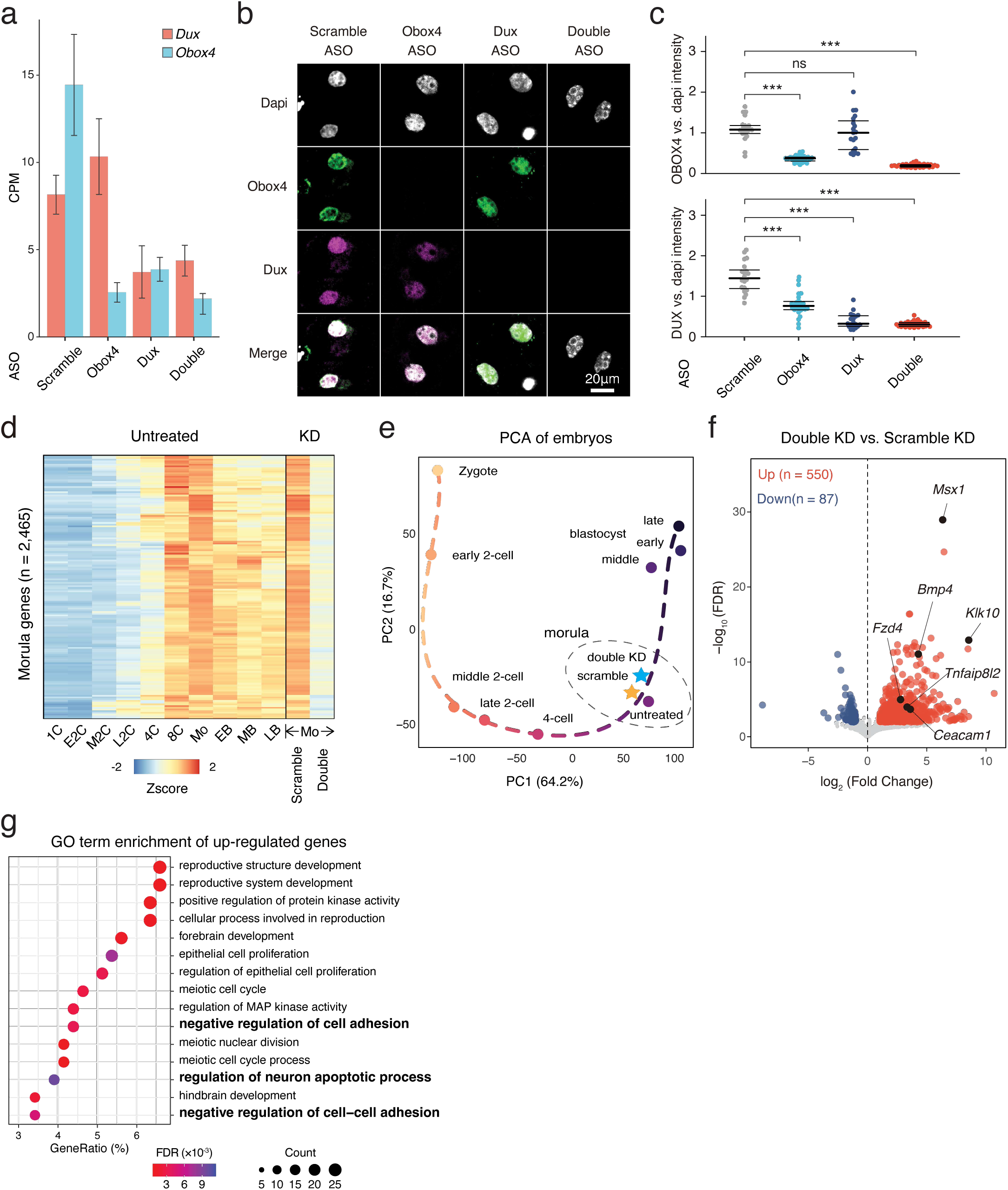
a) Transcript levels of *Dux* and *Obox4* in ASO-injected embryos, as measured using RNA-seq. CPM, counts per million. b) Representative images of immunofluorescence co-staining of OBOX4 and DUX in 2C embryos subjected to ASO-mediated knockdown. c) Dot plots showing the per-nucleus intensity of OBOX4 and DUX normalized to dapi. d) Heatmaps of the expression of morula-genes in preimplantation embryos and knockdown morulae. Genes appeared in both scRNA-seq cluster 6 and bulk RNA-seq cluster 3 in Supplementary Fig. 1a were designated as morula-genes. e) Transcriptome-based PCA of knockdown morulae and preimplantation embryos. f) Volcano plot of DEGs in double knockdown morulae comparing with scramble knockdown. Selected apoptosis associated genes are highlighted. g) The enrichment of gene ontology (GO) terms of up-regulated genes in double knockdown morulae.

**Supplementary Figure 11.**
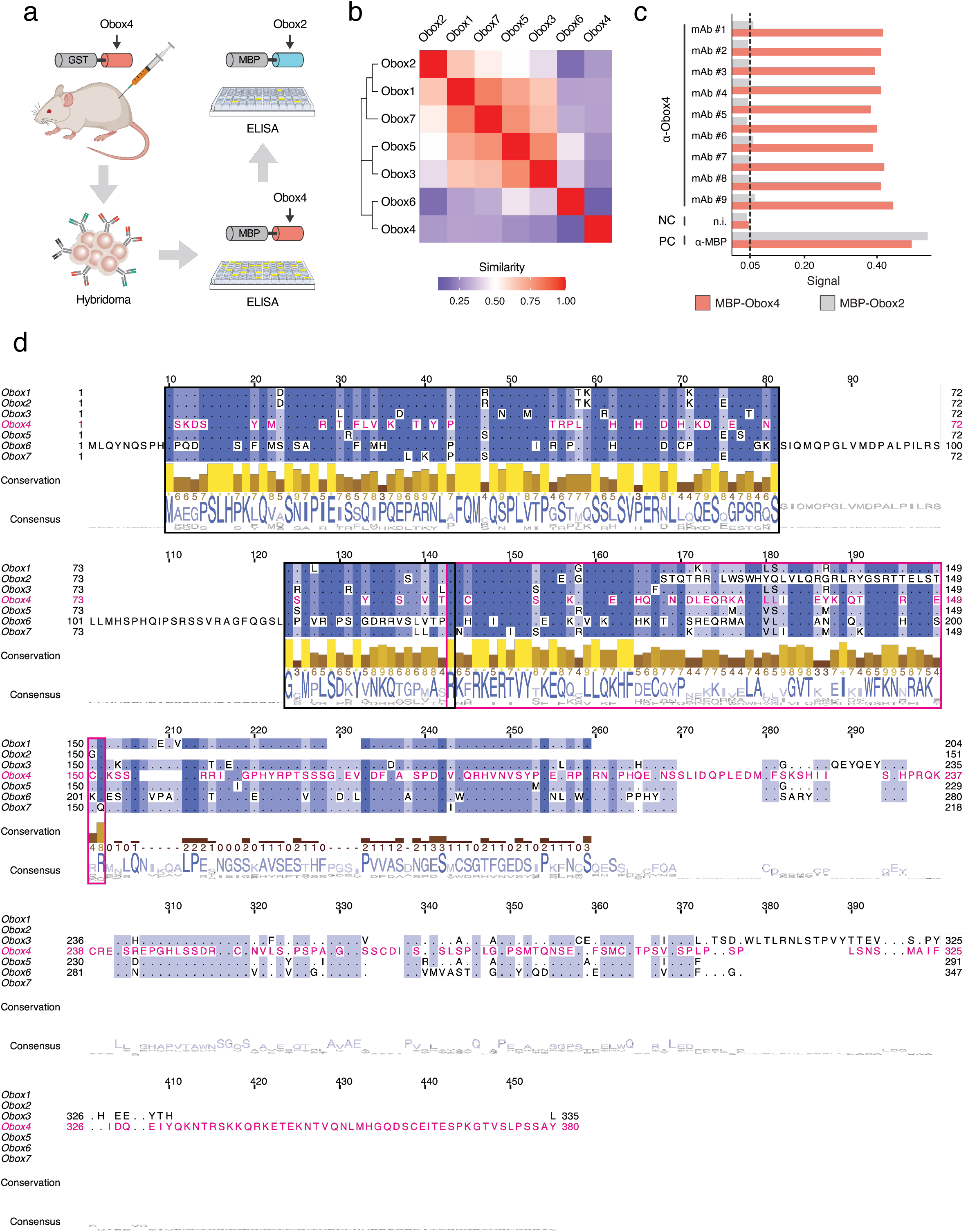
a) Schematic of the production of monoclonal antibodies. Mice were immunized with the GST-OBOX4 fusion protein. After fusing B-cells with myelomas, the antibodies produced by hybridomas were examined against MBP-OBOX4 fusion protein, followed by MBP-OBOX2 fusion protein, for identification of OBOX4-specific antibodies. b) Analysis of protein sequence similarity among the OBOX family members. The Smith–Waterman local alignment method was used to align the amino acid sequences, and the divergence among different sequences was represented in the form of a block substitution matrix. c) ELISA signal of anti-OBOX4 monoclonal antibodies against the MBP-OBOX4 and MBP-OBOX2 fusion proteins. Anti-OBOX4 monoclonal antibodies were not reactive to MBP-OBOX2. d) Multiple sequence alignment of protein sequence among the OBOX family members. The homeodomains are highlighted by magenta box. The peptide sequences used for antibody production are highlighted by black box.

## Methods

### Mice

Male/female BDF1 and female BALB/c mice were purchased from Japan SLC Inc. Mice were fed regular chow and housed in a controlled room under a 14 h/10 h light/dark cycle, at 22°C. All animal experiments were approved by the Animal Care and Use Committee of Keio University and the Animal Experimentation Committee at the RIKEN Tsukuba Institute and conducted in compliance with the Keio University Code of Research Ethics and the RIKEN’s guiding principles.

### Cell culture

All mouse mESC lines were cultured in Dulbecco’s Modified Eagle Medium (DMEM) (catalog no. 08488-55, Nacalai Tesque) supplemented with 10% fetal bovine serum (lot no. S10581S1820, Biowest), 1× GlutaMAX™ (catalog no. 35050061, Gibco), 1× penicillin-streptomycin (catalog no. 15140163, Gibco), 1× non-essential amino acids (catalog no. 11140050, Gibco), 1× sodium pyruvate (catalog no. S8636, Merck), 50 μM 2-mercaptoethanol (catalog no. 21985023, Gibco), 1 μM CHIR99021 (catalog no. 034-23103, Wako), 3 μM PD0325901 (catalog no. 168-25293, Wako), and mouse leukemia inhibitory factor produced in-house. The culture medium was supplemented with iMatrix-511 silk (recombinant human laminin-511 E8 fragment) (catalog no. 892021, Matrixome) at a concentration of 125 ng.cm^-2^ culture vessel area, before seeding mESCs into it. The cells were cultured in a humidified atmosphere containing 5% CO_2_ at 37°C, with the media changed every 48 h.

TBLCs were generated by culturing mESCs in an ordinary medium supplemented with 2.5 nM PLaB (catalog no. 16538, Cayman Chemicals). Cells were sub-cultured every 2–4 d, at a seeding density of 1×10^5^ cm^-2^ culture vessel area. After five rounds of sub-culture, the cells were deemed TBLCs.

SP2/O-Ag14 myeloma and primary clones of hybridomas were cultured in GIT medium (catalog no. 63725715, Wako) supplemented with recombinant human interleukin-6 (IL-6) (catalog no. 20006, PeproTech). Cells were cultured in a humidified atmosphere containing 5% CO_2_ at 37°C and sub-cultured every day, at a seeding density of 3×10^5^ mL^-1^. For monoclonal antibody production, hybridomas were cultured in hybridoma serum-free medium supplemented with IL-6. The cells were cultured in a humidified atmosphere containing 5% CO_2_ at 37°C, until they were over-confluent.

### Cell transfection

Plasmid transfections were performed using jetOPTIMUS^®^ DNA Transfection Reagent (catalog no. 101000025, Polyplus), according to the manufacturer’s instructions. Briefly, mESCs were seeded at a cell density of 8×10^4^ cm^-2^ culture vessel area. After 20 min, the jetOPTIMUS^®^ reagent and plasmids were diluted in jetOPTIMUS^®^ buffer, incubated for 10 min at room temperature, and then applied to the cell culture. After 48 h, the cells were collected for downstream experiments.

### cDNA synthesis and cloning

Total RNA was extracted from TBLCs using ISOGEN (catalog no. 311-02501, Nippon Gene), according to the manufacturer’s instructions. Briefly, 500 μL ISOGEN was added to 1.5×10^6^ freshly harvested TBLCs. After 5 min of incubation at room temperature, 100 μL of chloroform (catalog no. 03802606, Wako) was added to the cells and mixed by means of vigorous shaking. After 2 min of incubation at room temperature, the samples were centrifuged at 12,000 × *g* for 15 min at 4°C. The upper aqueous phase was then transferred to a new tube and mixed with 240 μL isopropanol (catalog no. 15-2320, Merck), to precipitate the RNA. After 5 min of incubation at room temperature, the samples were centrifuged at 12,000 × *g* for 15 min at 4°C. The supernatant was then discarded, following which the RNA pellets were washed with 70% ethanol (catalog no. 057-00456, Wako). The RNA pellets were then air-dried and dissolved in RNase-free water. The RNA solution was subjected to DNase treatment using TURBO DNA-*free*™ kit (catalog no. AM1907, Thermo Fisher), according to the manufacturer’s instructions, to remove genomic DNA carryover. RNA (1 μg) was reverse transcribed to cDNA using the Transcriptor First Strand cDNA Synthesis Kit (catalog no. 04379012001, Roche), according to the manufacturer’s instructions. PrimeSTAR^®^ Max DNA Polymerase (catalog no. R045A, TaKaRa) and ProFlex™ PCR System (catalog no. 4484073, Thermo Fisher) were used to amplify the sequence of interest from the cDNA. NEBuilder^®^ HiFi DNA Assembly Kit (catalog no. E2621L, NEB) was used to clone the sequence of interest into plasmid backbone.

### Immunofluorescence

For embryos, middle-2C-embryos were collected at 1.5 dpc. The zona pellucida was removed by treating the embryos with acidic Tyrode’s solution (catalog no. MR-0040-D, Merck). The embryos were fixed in 4% paraformaldehyde (catalog no. 09154-14, Nacalai Tesque) in PBS for 10 min. Following three washes in PBS, fixed embryos were permeabilized with 0.1% Triton™ X-100 (catalog no. 1610407, Bio-Rad) in PBS, for 20 min at room temperature. Following three washes in PBS, permeabilized embryos were blocked in 2% bovine serum albumin (BSA; catalog no. 011-27055, Wako) in PBS, for 20 min at room temperature. The blocked embryos were incubated with primary antibodies diluted in 2% BSA in PBS, at 4°C overnight. Following three washes in PBS, the embryos were incubated with 2% BSA in PBS containing 1:200 diluted 4′,6-diamidino-2-phenylindole (DAPI) (catalog no. 19178-91, Nacalai Tesque) and 1:500 diluted Alexa Fluor™-conjugated anti-mouse IgG1 secondary antibody (catalog no. A-21127, Thermo Fisher) or Alexa Fluor™-conjugated anti-mouse IgG2a secondary antibody (catalog no. A-21131, Thermo Fisher) and, for 1 h at room temperature. Following three washes in PBS, the embryos were transferred into liquid paraffin-covered PBS drops on a glass-bottom dish (catalog no. D11130H, Matsunami).

For culturing, cells seeded on glass bottom chamber slides (catalog no. SCS-N02, Matsunami) were fixed with 4% paraformaldehyde in PBS, for 10 min. After three washes in PBS, the fixed cells were permeabilized with 0.1% Triton™ X-100 in PBS, for 15 min at room temperature. Following three washes in PBS, permeabilized cells were incubated with primary antibodies diluted with 2% skim milk in PBS, for 30 min at room temperature. Following three washes in PBS, the cells were incubated with 2% BSA in PBS containing 1:500 diluted Alexa Fluor™-conjugated secondary antibody and 1:200 diluted DAPI, for 1 h at room temperature. Following three washes in PBS, the chamber was removed and the slides were mounted with ProLong™ Glass Antifade Mountant (catalog no. P36982, Thermo Fisher). Fluorescence images were taken using a confocal laser scanning microscope (catalog no. FV3000, Olympus).

### Western blot

Freshly harvested cells were re-suspended in phosphate-buffered saline (PBS) (catalog no. 14249-95, Nacalai Tesque) and lysed by means of sonication. Whole cell lysates were mixed with sample buffer containing reducing reagent (catalog no. 09499-14, Nacalai Tesque) and boiled at 95°C for 5 min. Protein samples were loaded on a 10% tris-glycine gel, run in AllView PAGE Buffer^®^ (catalog no. DS520, BioDynamics Laboratory), and then transferred to nitrocellulose membranes (catalog no. 10600003, Cytiva) using a Power Blotter System (catalog no. PB0012, Thermo Fisher). The membranes were blocked with PBST containing 2% skim milk (catalog no. 4902720131292, Morinaga), at room temperature for 15 min. Membranes were incubated with primary antibodies in PBST containing 2% skim milk, at room temperature for 30 min. After three washes with PBST, the membrane was incubated with secondary antibodies in PBST containing 2% skim milk, at room temperature for 15 min, with shaking. After three washes with PBST, the membrane was incubated with ECL reagents (catalog no. RPN2232, Cytiva) and exposed to an X-ray film (catalog no. 28906839, Cytiva) or detected using a digital chemiluminescence imager (catalog no. 17001402JA, Bio-Rad).

### qPCR

qPCR was performed using TB Green Fast qPCR Mix (catalog no. RR430A, TaKaRa) according to the manufacturer’s instructions. The qPCR reactions were carried out and the signals were detected using a real time PCR system (catalog no. TP950, TaKaRa). For RT-qPCR, first strand cDNA of mESCs total RNA was produced as described in methods **cDNA synthesis and cloning** section. Primers targeting MERVL Gag consensus sequence were used to detect MERVL transcript. Primers targeting beta actin mRNA were used as internal control in all qPCR assays performed in this study.

### smFISH

smFISH probes targeting MERVL were designed and synthesized by LGC Biosearch Technologies as previously described ^59^. smFISH followed by immunofluorescence staining was performed according to the manufacturer’s instructions. Briefly, Quasar 570 labeled probes were hybridized against MERVL RNA at 37°C for 16 h. Followed by immunofluorescence staining described in methods **Immunofluorescence** section, without blocking step to prevent RNase contamination. The slides were mounted with ProLong™ Glass Antifade Mountant (catalog no. P36982, Thermo Fisher). Fluorescence images were taken using a confocal laser scanning microscope (catalog no. FV3000, Olympus).

### Flow cytometry

The cells were digested with 0.25% trypsin-EDTA (catalog no. 25200072, Gibco), at 37°C for 5 min, and then re-suspended in FluoroBrite™ DMEM (catalog no. A1896701, Gibco). The cell suspension was filtered through a 35 μM cell strainer (catalog no. 352235, Falcon). The samples were analyzed using a cell sorter (catalog no. SH800Z, Sony). Data analysis was performed using the Sony SH800Z cell sorter software.

### Generation of transgenic mESC lines

The 2C::tdTomato reporter cell line was generated by transfection of EB3 mESCs (catalog no. AES0139, RIKEN BRC Cell Bank) with a linearized 2C::tdTomato reporter plasmid (catalog no. 40281, Addgene). After 48 h, the cells were subjected to 500 μg.mL^-1^ hygromycin (catalog no. 08906151, Wako) for 7 d. The selected cells were then seeded at a density of 2×10^2^ cm^-2^ in culture medium containing 250 μg.mL^-1^ hygromycin. Single-cell clones were picked and expanded after 7 d.

A tetracycycline-controlled transcription activation system and a piggyBac transposon system were used as previously described ^60^, to generate an *Obox4*-inducible cell line. Briefly, pPB-TRE-3×FLAG-Obox4, pPB-CAG-rtTA3G-IRES-Hygro, and pCMV-HyPBase-PGK-Puro plasmids were co-transfected into EB3 mESCs. After 48 h, the cells were subjected to selection with 500 μg.mL^-1^ hygromycin and 500 μg.mL^-1^ G418 (catalog no. 07805961, Wako), for 7 d. The selected cells were then seeded at a density of 2×10^2^ cm^-2^ in a culture medium containing 250 μg.mL^-1^ hygromycin and 250 μg.mL^-1^ G418. Single-cell clones were picked and expanded after 7 d.

### Generation of monoclonal antibodies

The anti-OBOX4, anti-DUX, and anti-GAG monoclonal antibodies were produced as previously described ^61^ (Supplementary Fig. 11a). Briefly, BALB/c mice were immunized by means of intraperitoneal injection of glutathione S-transferase-fused protein-of-interest (PoI). The mice were routinely immunized until their sera tested positive for the PoI-fused maltose binding protein (MPB), after which two boosting immunizations were performed. Following boosting, splenocytes from immunized mice were fused with SP2/O myeloma and cultured in hypoxanthine-aminopterin-thymidine medium for 10 d, to select for hybridomas. Hybridomas were subsequently screened using enzyme-linked immunosorbent assay against PoI-fused MBP. The positive polyclonal hybridomas were monoclonalized and expanded. For anti-OBOX4 hybridomas, additional screening using OBOX2-fused MBP was conducted. The sequence of OBOX4 was significantly divergent from those of other OBOX members, and the monoclonal antibodies were negative for cross-reactivity with OBOX2 (Supplementary Fig. 11b-c). The N-terminal 100 amino acids of OBOX4 and OBOX2 were used for immunization and cross-reactivity examination respectively (Supplementary Fig. 11d). The culture supernatants of monoclonal hybridomas were sterilized by passing them through a 0.22 μM pore size filter (catalog no. 430769, Corning) and used directly as an antibody solution in other assays.

### Generation of knockout mouse lines

Single guide RNAs (sgRNAs) designed on flanking and inside of the *Obox4* cluster were synthesized using a Precision gRNA Synthesis Kit (catalog no. A29377, Thermo Fisher) according to the manufacturer’s instructions. 150 ng.mL^-1^ sgRNA and 125 ng/μL Alt-R S.p. Cas9 Nuclease (catalog no. 1081058, Integrated DNA Technologies) diluted with Hepes-buffered KSOM were electroporated into BDF1 × B6 *in vitro* fertilized embryos with an electroporator (catalog no. NEPA21, NEPA GENE). Treated embryos were cultured in KSOM for 18 h and embryos that reached the two-cell stage were transferred into the oviducts of pseudopregnant ICR female mice on day 0.5. Pups were retrieved on day 19.5 and genotyped with specific PCR primers.

### Generation of knockout mESC lines

Plasmids encoding Cas9 and single guide RNA (sgRNA) flanking and inside of the *Obox4* cluster were constructed to perform knockout experiment. The *Obox4* knockout cell line was generated by means of co-transfection of EB3 mESCs with the pX330-Puro-Obox4KO-3, pX330-Puro-Obox4KO-4, pX330-Puro-Obox4KO-6, and pX330-Puro-Obox4KO-7 plasmids. After 18 h, the cells were subjected to selection with 1 μg.mL^-1^ puromycin (catalog no. 16023151, Wako) for 54 h. The selected cells were then seeded at a very low density in culture medium without antibiotics. Single-cell clones were picked and expanded after 7 d, and then subjected to genotyping.

Following a previously published protocol ^18^, plasmids encoding Cas9 and sgRNAs flanking the *Dux* cluster were constructed to perform knockout experiment. The *Dux* knockout cell line and the *Obox4*/*Dux* double knockout cell line were generated by means of co-transfection of EB3 mESCs and *Obox4* knockout cell lines with pX330-DuxKO-A and pX330-DuxKO-B plasmids. After 48 h, the cells were seeded at a very low density. Single-cell clones were picked and expanded after 7 d and then subjected to genotyping.

While mono-allelic *Obox4* deletion was detected in 49.1% (335/682) of the CRISPR-Cas9 edited clones, bi-allelic deletion was only detected in 0.44% (3/682) of the population, suggesting that removal of *Obox4* allele was associated with severe genetic toxicity. In contrast, 36.7% (47/128) mono-allelic and 4.7% (6/128) bi-allelic deletions were detected in *Dux* knockout clones.

### Genotyping and copy number examination

For mouse and culture cell, mouse right hindlimb toe or tail tip, or 1×10^6^ culture cells were incubated in 400 µL ProK solution containing 20 mM Tris-HCl pH = 7.9, 1 mM EDTA, 1% w/v sodium dodecyl sulfate (SDS), 150 mM NaCl, 20 mM trisodium citrate, and 80 µg recombinant proteinase K (catalog no. 161-28701, Wako) at 55°C for 2 h with vigorous shaking. The solution was extracted with 2 volumes of phenol/chloroform/isoamyl alcohol (25:24:1) (catalog no. 311-90151, Nippon Gene) twice. DNA in the aqueous phase was precipitated by adding 0.1 volume of 3 M sodium acetate and 2 volumes of 99.5% ethanol (catalog no. 057-00456, Wako), snap freeze in liquid nitrogen, and centrifuged at 15,000 × *g* for 12 min at 4°C. After discarding supernatant, the DNA pellets were washed by 200 µL 70% ethanol, air dried at room temperature, and dissolved in 220 µL TE solution (Tris-HCl pH = 7.9, 1 mM EDTA) supplemented with 1 µg RNase A (catalog no. 131-01461, Nippon Gene). The solution was incubated at 37°C for 1 h, then phenol/chloroform/isoamyl alcohol extracted and ethanol precipitated again as described above, to obtain highly purified genomic DNA. The DNA was dissolved in 200 µL TE solution.

For single blastomeres and abnormal embryos, whole genome amplifications were performed on individual embryos by using REPLI-g Advanced DNA Single Cell Kit (catalog no. 150363, Qiagen) according to the manufacturer’s instructions. For blastocysts, crude genomic DNA from single blastocysts was prepared according to the method described by Sakurai *et al*.^62^ with some modifications. Briefly, single blastocysts in 0.5 µL KSOM medium were transferred to the bottom of 0.1 mL PCR tubes, followed by addition of 10 µL blastocyst lysis buffer containing of 120 µg/mL recombinant proteinase K, 100mM Tris-HCl pH = 7.9, 100mM KCl, 0.45% NP-40, and 10 µg/mL yeast tRNA (catalog no. AM7119, Thermo Fisher). After brief vortex and pulse spin, the tubes were incubated at 55°C for 10 min followed by 95°C for 10 min.

For PCR based genotyping, 10 ng purified genomic DNA, 10ng whole genome amplified DNA, or 2µL of blastocyst lysate were used as template and amplified with target specific primers by KOD One^®^ PCR Master Mix (catalog no. KMM-101, TOYOBO) according to the manufacturer’s instructions. Primers targeting mouse *Tardbp* were used as internal control when whole genome amplification product or blastocyst lysate were used as multiplex PCR template. For copy number examination, 5 ng genomic DNA template was amplified as described in methods **qPCR** section.

### *In vitro* transcription

The *Obox4* coding sequence was codon-optimized using GeneArt Instant Designer (Thermo Fisher), to remove the ASO target motif and improve translation efficiency. Codon-optimized DNA was synthesized using Prime Gene Synthesis Services (Thermo Fisher). Codon-optimized *Obox4* mRNA was transcribed *in vitro* using the mMESSAGE mMACHINE™ T7 Transcription Kit (catalog no. AM1344, Thermo Fisher) and then polyadenylated using a poly(A) tailing kit (catalog no. AM1350, Thermo Fisher), according to the manufacturer’s instructions. The polyadenylated mRNA was purified using the RNeasy Mini Kit (catalog no. 74004, Qiagen), according to the manufacturer’s instructions, and dissolved in RNase-free water, at a concentration of 100 ng.μL^-1^.

### Pronuclear injection of mouse embryos

Eight-week-old BDF1 female mice were injected with 150 µL of CARD HyperOva (catalog no. KYD-010-EX, Kyudo Co., Ltd.). After 48 hours, the female mice were injected with 5 IU human chorionic gonadotropin (hCG) (catalog no. GONATROPIN, ASKA Animal Health Co. Ltd.) and mated with BDF1 male mice. After 22 h, the PN3 zygotes were collected. Cumulus cells were removed by briefly culturing the zygotes in potassium simplex optimization medium (KSOM) (catalog no. MR-101-D, Merck) supplemented with 0.3 µg.µL^-1^ hyaluronidase (catalog no. H4272, Merck). Embryos were cultured in KSOM medium drops covered with liquid paraffin (catalog no. 26137-85, Nacalai Tesque), in a humidified atmosphere with 5% CO2, at 37°C. After 4 h, PBS containing either 40 µM scramble ASO, 20 µM anti-*Dux* ASO, 20 µM anti-*Obox4* ASO, 40 µM equimolar anti-*Dux*/anti-*Obox4* ASO mixture, or 40 µM equimolar anti-*Dux*/anti-*Obox4* ASO mixture with 100 ng.μL^-1^ codon-optimized *Obox4* mRNA was microinjected into the male pronuclei of zygotes using a microinjector (catalog no. 5252000013, Eppendorf). For developmental monitoring, embryos were cultured for 4 d after microinjection and assessed for developmental stage at 18 h, 42 h, 66 h, and 90 h after microinjection, which corresponded to 1.5 dpc, 2.5 dpc, 3.5 dpc, and 4.5 dpc, respectively.

### Somatic cell nuclear transfer (SCNT)

SCNT was performed as described previously ^63^ using wild-type and knockout mESC lines as nuclear donor cells. They were cultured at a high density until they reached confluency for about four to six days before SCNT. BDF1 female mice were superovulated by the injection of 7.5 IU of equine chorionic gonadotropin (eCG, ZENOAQ) and 7.5 IU of hCG (ASKA Pharmaceutical Co., Ltd.) at a 48 h interval. At 15 h after hCG injection, cumulus–oocyte complexes were collected from the oviducts. After removal of cumulus cells by 0.1% bovine testicular hyaluronidase (catlog no. 385931, Calbiochem), oocytes were enucleated in Hepes-buffered KSOM containing 7.5 µg/mL cytochalasin B. After culture in KSOM for at least 1 h, the enucleated oocytes were injected with donor mESCs using a Piezo-driven micromanipulator (catalog no. PMM-150FU, Primetech). After culture in KSOM for about 1 h, injected oocytes were cultured in Ca^2+^-free KSOM containing 2.5 mM SrCl2, 5 μM latrunculin A (LatA) (catalog no. L5163, Merck) with 50 nM trichostatin A (TSA) (catalog no. T8552, Merck) for 1 h. Then, they were cultured in KSOM containing 5 µM LatA and 50 nM TSA for 7 h. After washing, the SCNT embryos were cultured in KSOM under 5% CO2 in air at 37°C for 96 h. The embryos that reached 2-cells at 24 h after activation were considered successfully cloned with cell cycle-matched mESCs (G0/G1 phase).

### scRNA-seq analysis

Raw scRNA-seq data was obtained from the dataset of Deng et al. (GSE45719). Quality control and adapter trimming was done using fastp ^64^ v0.23.2. Quality-controlled reads were aligned to the GRCm38.p6 reference genome using STAR ^65^ v2.7.9a, with default arguments. Reads were counted against GRCm38.p6 comprehensive gene annotation ^66^ and mm10 repeats from the University of California, Santa Cruz (UCSC) RepeatMasker using Subread ^67^ v2.0.1 featureCounts function, and multi-mapping reads were discarded for non-TE features and counted fractionally for TEs. Seurat ^68^ v4.1.0 was used to process the read counts of scRNA-seq. Cells with greater than 7.5% mitochondrial reads or less than 14,000 annotated features were discarded. Expression levels were log-normalized. The expression profiles of homeobox genes listed in HomeoDB2 ^69^ were clustered into 11 *k*-means after *z*-score transformation.

### Ribo-seq analysis

Raw Ribo-seq data was obtained from the dataset of Xiong et al. (GSE165782). Reads were quality-controlled, aligned, and counted, as described above. Reads aligned to all protein coding *Obox4* loci were added up to represent translation level of OBOX4.

### Bulk RNA-seq

For embryos, 30 middle-2C-embryos were collected 20 h after microinjection in each independent biological replicate. The zona pellucida was removed by treating embryos with acidic Tyrode’s solution. Libraries were constructed using the SMART-Seq^®^ Stranded Kit (catalog no. 634442, TaKaRa) and indexed using the SMARTer^®^ RNA Unique Dual Index Kit (catalog no. 634451, TaKaRa), according to the manufacturer’s instructions.

For cell culture, total RNA was prepared using the RNeasy Kit, according to the manufacturer’s instructions. The RNA solution was subjected to DNase treatment to remove genomic DNA carryover. Using 1 µg total RNA per sample, libraries were constructed with the NEBNext^®^ Ultra™ II Directional RNA Library Prep Kit for Illumina (catalog no. E7760L, NEB) and indexed with NEBNext^®^ Multiplex Oligos for Illumina (catalog no. E6440S, NEB), according to the manufacturer’s instructions.

The libraries were quantified with a 2100 Bioanalyzer (catalog no. G2939BA, Agilent) using a High Sensitivity DNA Kit (catalog no. 5067-4626, Agilent). Quantified libraries were pooled and sequenced using an Illumina NovaSeq 6000 System in 150 bp paired-end mode (Illumina). Base calling and de-multiplexing were performed using the bcl2fastq2 (Illumina) v2.20. De-multiplexed reads were quality-controlled, aligned, and counted, as described above. DESeq2 ^70^ v1.32.0 was used to perform differential expression analysis. Raw RNA-seq reads for TBLCs were downloaded from the dataset of Shen et al. (GSE168728). Raw RNA-seq reads of *Dux* knockout embryos were downloaded from the datasets of Chen & Zhang and De Iaco et al. (GSE121746 and GSE141321 respectively). Raw RNA-seq reads of preimplantation embryos were downloaded from the datasets of Wu et al. (GSE66390). Raw RNA-seq reads of 2C-like mESCs were downloaded from the datasets of Zhu et al. (GSE159623). The published data were analyzed using the same method.

### CUT&RUN-seq

Cultured cells (1×10^5^) freshly prepared by trypsin-EDTA digestion were used to perform CUT&RUN with the CUT&RUN Assay Kit (catalog no. 86652, Cell Signaling Technology), according to the manufacturer’s instructions. Enriched DNA was subjected to library construction using the NEBNext^®^ Ultra™ II DNA Library Prep Kit for Illumina (catalog no. E7645L, NEB), with the adapter ligation step performed at 50°C instead of 65°C, to prevent the denaturation of small DNA inserts. The libraries were indexed using NEBNext^®^ Multiplex Oligos for Illumina, according to the manufacturer’s instructions.

The libraries were quantified, pooled, sequenced, de-multiplexed, and quality controlled, as described above. Processed reads were aligned to the GRCm38.p6 reference genome using Bowtie2 ^71^ v2.4.1, with default settings. Peaks were called in each biological replicate using the MACS3 ^72^ v3.0.0a6 callpeak function (-f BAMPE). Alignment tracks were first generated using deepTools ^73^ v3.5.1 bamCoverage function (--binSize 10 --normalizeUsing CPM --smoothLength 30), and then normalized by subtracting the signal from non-immune IgG and wildtype mESCs using the deepTools bamCompare function (--scaleFactorsMethod None --operation subtract --binSize 10 --smoothLength 30). ChIPseeker ^74^ v1.28.3 was used to annotate the peaks. The MEME suite ^75^ v5.4.1 was used to identify the binding motifs. Heatmaps were generated using deepTools computeMatrix and plotHeatmap functions. Visualization of genomic tracks was performed using trackplot ^76^ v1.3.10. Raw ChIP-seq reads of DUXs in mESCs were downloaded from Hendrickson et al. (GSE85632). The published data were analyzed using the same method.

### Quantification and statistical analysis

Descriptive and comparative statistics were employed in the manuscript as described in the figure legends with the number of replicates indicated. Significance is defined as a *p-value* less than 0.05 indicated with asterisk (* *p-value* < 0.05, ** *p-value* < 0.01, *** *p-value* < 0.001). Error bar represents the standard deviation (SD) of the mean of the replicates.

## Data and code availability

The RNA-seq and CUT&RUN-seq data generated in this study have been deposited at NCBI Gene Expression Omnibus (GEO) database under the accession code GSE196671.

## Acknowledgements

We thank all the members of the Siomi Laboratory for their discussions and comments on this work. We thank Takehiko Yokomizo (Department of Biochemistry, Juntendo University) for providing the anti-DYKDDDDK monoclonal antibody (clone 2H8). We also thank Daisuke Motooka (Research Institute for Microbial Diseases, Osaka University) for generating the sequencing data. We are grateful to Azusa Inoue (Center for Integrative Medical Sciences, RIKEN) and Katsuhiko Hayashi (Department of Genome Biology, Osaka University) for their comments on this manuscript. This work was supported by the MEXT Grant-in-Aid for Scientific Research in Innovative Areas (19H05753 to H.S. and 19H05758 to A.O.), AMED project to elucidate and control mechanisms of aging and longevity (1005442 to H.S.), JSPS Grant-in-Aid for Scientific Research KAKENHI (20K21507 to K.M. and 22H02534 to K.I.), Mochida Memorial Foundation Research Grant to K.M., Sumitomo Foundation Research Grant to K.M., Keio University Doctorate Student Grant-in-Aid Program to Y.G., and the JST Doctoral Program Student Support Fellowship to Y.G.

## Author contributions

H.S. and Y.G. conceived and designed the project; Y.G., T.K., K.M., T.D.L., A.S., H.I., N.O., S.M., M.S., and A. O. performed the experiments; Y.G. and T.K. analyzed the data; H.I. developed the ASO-mediated knockdown experiments; H.S. and K.M. supervised and coordinated the experiments; Y.G. and H.S. wrote the paper with input from all authors. All authors reviewed and approved the final version of the manuscript.

## Declaration of interests

The authors declare no competing interests.

## Notes

### Competing Interest Statement

The authors have declared no competing interest.

### Summary of Updates

Produced Dux and Obox4 single/double knockout mice. Added fertility data, preimplantation development monitoring, transcriptome profiling analysis of single and double KO embryos. Removed histone modification analysis in TBLCs for better coherence of the manuscript.

## References

1. Lee, M.T., Bonneau, A.R. & Giraldez, A.J. Zygotic genome activation during the maternal-to-zygotic transition. Annual review of cell and developmental biology 30, 581–613 (2014).

2. Tarkowski, A.K. Experiments on the development of isolated blastomeres of mouse eggs. Nature 184, 1286–1287 (1959).

3. Tarkowski, A.K. Experimental studies on regulation in the development of isolated blastomeres of mouse eggs; Badania eksperymentalne nad rozwojem izolowanych blastomerów jaj myszy. Acta Theriologica 3, 191–267 (1959).

4. Suwińska, A., Czołowska, R., Ożdżeński, W. & Tarkowski, A.K. Blastomeres of the mouse embryo lose totipotency after the fifth cleavage division: expression of Cdx2 and Oct4 and developmental potential of inner and outer blastomeres of 16-and 32-cell embryos. Developmental biology 322, 133–144 (2008).

5. Svoboda, P. et al. RNAi and expression of retrotransposons MuERV-L and IAP in preimplantation mouse embryos. Developmental biology 269, 276–285 (2004).

6. McGinnis, W., Levine, M.S., Hafen, E., Kuroiwa, A. & Gehring, W.J. A conserved DNA sequence in homoeotic genes of the Drosophila Antennapedia and bithorax complexes. Nature 308, 428–433 (1984).

7. Carrasco, A.E., McGinnis, W., Gehring, W.J. & De Robertis, E.M. Cloning of an X. laevis gene expressed during early embryogenesis coding for a peptide region homologous to Drosophila homeotic genes. Cell 37, 409–414 (1984).

8. McGinnis, W., Hart, C.P., Gehring, W.J. & Ruddle, F.H. Molecular cloning and chromosome mapping of a mouse DNA sequence homologous to homeotic genes of drosophila. Cell 38, 675–680 (1984).

9. Scott, M.P. & Weiner, A.J. Structural relationships among genes that control development: sequence homology between the Antennapedia, Ultrabithorax, and fushi tarazu loci of Drosophila. Proceedings of the National Academy of Sciences 81, 4115–4119 (1984).

10. Gehring, W.J. & Hiromi, Y. Homeotic genes and the homeobox. Annual review of genetics 20, 147–173 (1986).

11. Gehring, W.J. Homeo boxes in the study of development. Science 236, 1245–1252 (1987).

12. Lewin, T.D., Royall, A.H. & Holland, P.W. Dynamic Molecular Evolution of Mammalian Homeobox Genes: Duplication, Loss, Divergence and Gene Conversion Sculpt PRD Class Repertoires. Journal of Molecular Evolution, 1–19 (2021).

13. Töhönen, V. et al. Novel PRD-like homeodomain transcription factors and retrotransposon elements in early human development. Nature communications 6, 8207 (2015).

14. Madissoon, E. et al. Characterization and target genes of nine human PRD-like homeobox domain genes expressed exclusively in early embryos. Scientific reports 6, 1–14 (2016).

15. De Iaco, A. et al. DUX-family transcription factors regulate zygotic genome activation in placental mammals. Nature genetics 49, 941–945 (2017).

16. Hendrickson, P.G. et al. Conserved roles of mouse DUX and human DUX4 in activating cleavage-stage genes and MERVL/HERVL retrotransposons. Nature genetics 49, 925–934 (2017).

17. Whiddon, J.L., Langford, A.T., Wong, C.-J., Zhong, J.W. & Tapscott, S.J. Conservation and innovation in the DUX4-family gene network. Nature genetics 49, 935–940 (2017).

18. Chen, Z. & Zhang, Y. Loss of DUX causes minor defects in zygotic genome activation and is compatible with mouse development. Nature genetics 51, 947–951 (2019).

19. Guo, M. et al. Precise temporal regulation of Dux is important for embryo development. Cell research 29, 956–959 (2019).

20. De Iaco, A., Verp, S., Offner, S., Grun, D. & Trono, D. DUX is a non-essential synchronizer of zygotic genome activation. Development 147, dev177725 (2020).

21. Bosnakovski, D., Gearhart, M.D., Ho Choi, S. & Kyba, M. Dux facilitates post-implantation development, but is not essential for zygotic genome activation. Biology of reproduction 104, 83–93 (2021).

22. Shen, H. et al. Mouse totipotent stem cells captured and maintained through spliceosomal repression. Cell 184, 2843–2859. e20 (2021).

23. Wagner, A. Genetic redundancy caused by gene duplications and its evolution in networks of transcriptional regulators. Biological cybernetics 74, 557–567 (1996).

24. Deng, Q., Ramsköld, D., Reinius, B. & Sandberg, R. Single-cell RNA-seq reveals dynamic, random monoallelic gene expression in mammalian cells. Science 343, 193–196 (2014).

25. Howe, K.L. et al. Ensembl 2021. Nucleic Acids Research 49, D884–D891 (2020).

26. Xiong, Z. et al. Ultrasensitive Ribo-seq reveals translational landscapes during mammalian oocyte-to-embryo transition and pre-implantation development. Nature Cell Biology 24, 968–980 (2022).

27. Macfarlan, T.S. et al. Endogenous retroviruses and neighboring genes are coordinately repressed by LSD1/KDM1A. Genes & development 25, 594–607 (2011).

28. Macfarlan, T.S. et al. Embryonic stem cell potency fluctuates with endogenous retrovirus activity. Nature 487, 57–63 (2012).

29. Wu, J. et al. The landscape of accessible chromatin in mammalian preimplantation embryos. Nature 534, 652–657 (2016).

30. Zhu, Y. et al. Relaxed 3D genome conformation facilitates the pluripotent to totipotent-like state transition in embryonic stem cells. Nucleic acids research 49, 12167–12177 (2021).

31. Skene, P.J. & Henikoff, S. An efficient targeted nuclease strategy for high-resolution mapping of DNA binding sites. Elife 6, e21856 (2017).

32. Sasaki, F. et al. A high-affinity monoclonal antibody against the FLAG tag useful for G-protein-coupled receptor study. Analytical biochemistry 425, 157–165 (2012).

33. Markoulaki, S., Meissner, A. & Jaenisch, R. Somatic cell nuclear transfer and derivation of embryonic stem cells in the mouse. Methods 45, 101–114 (2008).

34. Ji, S. et al. OBOX regulates mouse zygotic genome activation and early development. Nature 620, 1047–1053 (2023).

35. Rajkovic, A., Yan, C., Yan, W., Klysik, M. & Matzuk, M.M. Obox, a family of homeobox genes preferentially expressed in germ cells. Genomics 79, 711–717 (2002).

36. Ge, S.X. Exploratory bioinformatics investigation reveals importance of “junk” DNA in early embryo development. BMC genomics 18, 1–19 (2017).

37. Zhong, Y.-f. & Holland, P.W. The dynamics of vertebrate homeobox gene evolution: gain and loss of genes in mouse and human lineages. BMC evolutionary biology 11, 1–13 (2011).

38. Wilming, L.G., Boychenko, V. & Harrow, J.L. Comprehensive comparative homeobox gene annotation in human and mouse. Database 2015(2015).

39. Lee, H.-S. & Lee, K.-A. Characterization and functional analysis of Obox4 during oocyte maturation by RNA interference. Clinical and Experimental Reproductive Medicine 34, 293–303 (2007).

40. Lee, H.S. et al. Obox4 critically regulates cAMP-dependent meiotic arrest and MI-MII transition in oocytes. The FASEB Journal 24, 2314–2324 (2010).

41. Lee, K.-A., Lee, H.-S., Kim, E.-Y., Kim, K.-H. & Kim, Y. Oocyte-specific homeobox 4 (Obox4) is a key regulatory component of the cAMP-dependent meiotic arrest during in vitro oocyte maturation. Fertility and Sterility 90, S108 (2008).

42. Lee, H.-S., Kim, E.-Y. & Lee, K.-A. Changes in gene expression associated with oocyte meiosis after Obox4 RNAi. Clinical and experimental reproductive medicine 38, 68 (2011).

43. Lee, H.-S. et al. Obox4-silencing-activated STAT3 and MPF/MAPK signaling accelerate nuclear membrane breakdown in mouse oocytes. Reproduction 151, 369–378 (2016).

44. Kim, H.-M. et al. Obox4 regulates the expression of histone family genes and promotes differentiation of mouse embryonic stem cells. FEBS letters 584, 605–611 (2010).

45. Yang, L. et al. Transient Dux expression facilitates nuclear transfer and induced pluripotent stem cell reprogramming. EMBO reports 21, e50054 (2020).

46. Leidenroth, A. & Hewitt, J.E. A family history of DUX4: phylogenetic analysis of DUXA, B, C and Duxbl reveals the ancestral DUX gene. BMC evolutionary biology 10, 1–13 (2010).

47. Maeso, I. et al. Evolutionary origin and functional divergence of totipotent cell homeobox genes in eutherian mammals. BMC biology 14, 1–14 (2016).

48. Zou, Z. et al. Translatome and transcriptome co-profiling reveals a role of TPRXs in human zygotic genome activation. *Science*, eabo7923 (2022).

49. Nowak, M.A., Boerlijst, M.C., Cooke, J. & Smith, J.M. Evolution of genetic redundancy. Nature 388, 167–171 (1997).

50. Vavouri, T., Semple, J.I. & Lehner, B. Widespread conservation of genetic redundancy during a billion years of eukaryotic evolution. Trends in genetics 24, 485–488 (2008).

51. Hackett, J.A. & Surani, M.A. Regulatory principles of pluripotency: from the ground state up. Cell stem cell 15, 416–430 (2014).

52. Sakashita, A. et al. Endogenous retroviruses drive species-specific germline transcriptomes in mammals. Nature structural & molecular biology 27, 967–977 (2020).

53. Bachtrog, D. Y-chromosome evolution: emerging insights into processes of Y-chromosome degeneration. Nature Reviews Genetics 14, 113–124 (2013).

54. Zaret, K.S. Pioneer transcription factors initiating gene network changes. Annual review of genetics 54, 367 (2020).

55. Eckersley-Maslin, M.A., Alda-Catalinas, C. & Reik, W. Dynamics of the epigenetic landscape during the maternal-to-zygotic transition. Nature Reviews Molecular Cell Biology 19, 436–450 (2018).

56. Gassler, J. et al. Zygotic genome activation by the totipotency pioneer factor Nr5a2. Science 378, 1305–1315 (2022).

57. Lai, F. et al. NR5A2 connects zygotic genome activation to the first lineage segregation in totipotent embryos. Cell Research, 1–15 (2023).

58. Grow, E.J. et al. p53 convergently activates Dux/DUX4 in embryonic stem cells and in facioscapulohumeral muscular dystrophy cell models. Nature genetics 53, 1207–1220 (2021).

59. Sakashita, A. et al. Transcription of MERVL retrotransposons is required for preimplantation embryo development. Nature genetics 55, 484–495 (2023).

60. Takeuchi, C., Murano, K., Ishikawa, M., Okano, H. & Iwasaki, Y.W. Generation of Stable Drosophila Ovarian Somatic Cell Lines Using the piggyBac System. in piRNA: Methods and Protocols 143–153 (Springer, 2022).

61. Guo, Y. et al. Potent mouse monoclonal antibodies that block SARS-CoV-2 infection. Journal of Biological Chemistry 296(2021).

62. Sakurai, T., Watanabe, S., Kamiyoshi, A., Sato, M. & Shindo, T. A single blastocyst assay optimized for detecting CRISPR/Cas9 system-induced indel mutations in mice. BMC biotechnology 14, 1–11 (2014).

63. Inoue, K. et al. Loss of H3K27me3 imprinting in the Sfmbt2 miRNA cluster causes enlargement of cloned mouse placentas. Nature communications 11, 1–12 (2020).

64. Chen, S., Zhou, Y., Chen, Y. & Gu, J. fastp: an ultra-fast all-in-one FASTQ preprocessor. Bioinformatics 34, i884–i890 (2018).

65. Dobin, A. et al. STAR: ultrafast universal RNA-seq aligner. Bioinformatics 29, 15–21 (2013).

66. Frankish, A. et al. GENCODE 2021. Nucleic Acids Research 49, D916–D923 (2020).

67. Liao, Y., Smyth, G.K. & Shi, W. featureCounts: an efficient general purpose program for assigning sequence reads to genomic features. Bioinformatics 30, 923–930 (2014).

68. Hao, Y. et al. Integrated analysis of multimodal single-cell data. Cell (2021).

69. Zhong, Y.f. & Holland, P.W. HomeoDB2: functional expansion of a comparative homeobox gene database for evolutionary developmental biology. Evolution & development 13, 567–568 (2011).

70. Love, M.I., Huber, W. & Anders, S. Moderated estimation of fold change and dispersion for RNA-seq data with DESeq2. Genome biology 15, 1–21 (2014).

71. Langmead, B. & Salzberg, S.L. Fast gapped-read alignment with Bowtie 2. Nature methods 9, 357–359 (2012).

72. Zhang, Y. et al. Model-based analysis of ChIP-Seq (MACS). Genome biology 9, 1–9 (2008).

73. Ramírez, F. et al. deepTools2: a next generation web server for deep-sequencing data analysis. Nucleic acids research 44, W160–W165 (2016).

74. Yu, G., Wang, L.-G. & He, Q.-Y. ChIPseeker: an R/Bioconductor package for ChIP peak annotation, comparison and visualization. Bioinformatics 31, 2382–2383 (2015).

75. Bailey, T.L. et al. MEME SUITE: tools for motif discovery and searching. Nucleic acids research 37, W202–W208 (2009).

76. Pohl, A. & Beato, M. bwtool: a tool for bigWig files. Bioinformatics 30, 1618–1619 (2014).

