## Supplementary Table 8 for "*Obox4* promotes zygotic genome activation upon loss of *Dux*"

**Supplementary Table 7. Primers used in this study**

| **Name** | **Sequences (5’ to 3’)**  **Comment** |
| --- | --- |
| Dux_clone_Fwd | GCATATGCAGGAATTC**ATGGCAGAAGCTGGCAGCCCTGTTGG** |
| Dux_clone_Rev | GCTTGATATCGAATTC**TCAGAGCATATCTAGAAGAGTCTGATATTC** |
| Obox4_clone_Fwd | GCATATGCAGGAATTC**ATGTCCAAAGATTCCTCCTTGCATCC** |
| Obox4_clone_Rev | GCTTGATATCGAATTC**GCGCTATAGAGACATCATGGCAT** |
| Emx2_clone_Fwd | GCATATGCAGGAATTC**ATGTTTCAGCCGGCGCCC** |
| Emx2_clone_Rev | GCTTGATATCGAATTC**TTAATCGTCTGAGGTCACATCTATTTCCTC** |
| Hoxd13_clone_Fwd | GCATATGCAGGAATTC**ATGAGCCGCTCGGGGACTTG** |
| Hoxd13_clone_Rev | GCTTGATATCGAATTC**TCAGGAGACAGTGTCTTTGAGCTTGG** |
| Actb_qPCR_Fwd | **gtgtgacgttgacatccgtaaagac** |
| Actb_qPCR_Rev | **gatcttcatggtgctaggagcca** |
| MERVL_Gag_qPCR_Fwd | **aaacaggctcctagaggggag** |
| MERVL_Gag­_qPCR_Rev | **tccacccttatcccacaccc** |
| Dux_pGEX_Fwd | TGGGATCCCCGAATTC**TGGGGTAGAAATCCTGGCAGC** |
| Dux_pGEX_Rev | GTCGACCCGGGAATTCTCA**TGCTTGAGTGCGGGCATCTTC** |
| Dux_pMAL_Fwd | CGGCACACTACGTAGAATTC**TGGGGTAGAAATCCTGGCAGC** |
| Dux_pMAL_Rev | CTAGAGGATCCGAATTCTCA**TGCTTGAGTGCGGGCATCTTC** |
| Obox4_pGEX_Fwd | TGGGATCCCCGAATTC**ATGTCCAAAGATTCCTCCTTGCATCC** |
| Obox4_pGEX_Rev | GTCGACCCGGGAATTCTCA**CACGGTGCGTTCCTTTCGAC** |
| Obox4_pMAL_Fwd | CGGCACACTACGTAGAATTC**ATGTCCAAAGATTCCTCCTTGCATCC** |
| Obox4_pMAL_Rev | CTAGAGGATCCGAATTCTCA**CACGGTGCGTTCCTTTCGAC** |
| Obox2_pMAL_Fwd | CGGCACACTACGTAGAATTC**ATGGCGGAAGGTCCCTCCTTGC** |
| Obox2_pMAL_Rev | CTAGAGGATCCGAATTCTCA**CACAGTTCGTTCTTTTCGAAACTTTCTTG** |
| Gag_pGEX_Fwd | TGGGATCCCCGAATTC**AATCTTTTAAAATACTGGAATTGGCTTGTTGATCC** |
| Gag_pGEX_Rev | GTCGACCCGGGAATTCTCA**TACTTCCTCAGGTGGGGCAAACC** |
| Gag_pMAL_Fwd | CGGCACACTACGTAGAATTC**AATCTTTTAAAATACTGGAATTGGCTTGTTGATC** |
| Gag_pMAL_Rev | CTAGAGGATCCGAATTCTCA**TACTTCCTCAGGTGGGGCAAACC** |
| Dux_KO_A_Fwd | CACC**AAGGCACACAGCCGCTTGCT** |
| Dux_KO_A_Rev | AAAC**AGCAAGCGGCTGTGTGCCTT** |
| Dux_KO_B_Fwd | CACC**GACTTTCCCCACTAGTGGCT** |
| Dux_KO_B_Rev | AAAC**AGCCACTAGTGGGGAAAGTC** |
| Obox4_KO_A_Fwd | cacc**GGAAGAGCTGTACCCCTAGA** |
| Obox4_KO_A_Rev | aaac**TCTAGGGGTACAGCTCTTCC** |
| Obox4_KO_B_Fwd | cacc**GAGAAGGAACAATGTGCCTA** |
| Obox4_KO_B_Rev | aaac**TAGGCACATTGTTCCTTCTC** |
| Dux_KO_genotyping_A | **AACTAGGGCATTTCTTACCCTGCTTGCC** |
| Dux_KO_genotyping_B | **ACCACTGCACATACATTAGAGACGTTTGG** |
| Dux_KO_genotyping_C | **CTGATAGACTGTGGGTTTCTTAGTGGC** |
| Dux_KO_genotyping_D | **ATAAACTAGAGGACAGAGCAAGTGGCC** |
| Dux_KO_genotyping_E | **ACAACACCTGATCTGCCTCTCACTCC** |
| Dux_KO_genotyping_F | **CAAACCCAACAAAGCCAACCCACC** |
| Obox4_KO_genotyping_A | **GAATCCTGATAACTGCTATCTAAGAGTCC** |
| Obox4_KO_genotyping_B | **AGGAACTTCACCGCCATAGCTGG** |
| Obox4_KO_genotyping_C | **CCTTCATGGCTCATTGGGTATTG** |
| Obox4_KO_genotyping_D | **CCATCTTTCACGATCAGAATGGC** |
| Obox4_KO_genotyping_E | **CCTCTAGTGACCCCAACACGTC** |
| Obox4_KO_genotyping_F | **GCCACTGCTGGAAGTAGGCCTA** |
| Tardbp_qPCR_Fwd | **gtggagctggcttgggaaataac** |
| Tardbp_qPCR_Rev | **cagaattagaaccactgtaggaattatttccag** |
