## Supplementary Table 9 for "*Obox4* promotes zygotic genome activation upon loss of *Dux*"

**Supplementary Table 8. Antisense oligonucleotides sequences used in this study**

| **Name** | **Sequences (5’ to 3’)**  **Comment** |
| --- | --- |
| ASO_scramble | /52MOEr**A**/*/i2MOEr**G**/*/i2MOEr**C**/*/i2MOEr**G**/*/i2MOEr**C**/***G*****G*****G*****T*****A*****T*****T*****G*****A*****A***/i2MOEr**C**/*/i2MOEr**C**/*/i2MOEr**A**/*/i2MOEr**G**/*/32MOEr**G**/ |
| ASO_Dux | /52MOEr**G**/*/i2MOEr**A**/*/i2MOEr**T**/*/i2MOEr**T**/*/i2MOEr**C**/***C*****T*****G*****C*****G*****G*****T*****T*****C*****T***/i2MOEr**G**/*/i2MOEr**A**/*/i2MOEr**A**/*/i2MOEr**A**/*/32MOEr**C**/ |
| ASO_Obox4 | /52MOEr**A**/*/i2MOEr**C**/*/i2MOEr**T**/*/i2MOEr**T**/*/i2MOEr**G**/***A*****T*****T*****G*****C*****A*****G*****T*****G*****G***/i2MOEr**A**/*/i2MOEr**C**/*/i2MOEr**G**/*/i2MOEr**T**/*/32MOEr**G**/ |
| **Modification** | **Chemical group** |
| 52MOEr | 5’ 2-MethoxyEthoxy |
| 32MOEr | 3’ 2-MethoxyEthoxy |
| i2MOEr | Internal 2-MethoxyEthoxy |
| * | Phosphorothioate Bond |
