## Supplementary Table 10 for "*Obox4* promotes zygotic genome activation upon loss of *Dux*"

**Supplementary Table 9. sgRNA sequences used in this study**

| **Name** | **Sequences (5’ to 3’)**  **Comment** | **PAM** |
| --- | --- | --- |
| Dux_KO_A | AAGGCACACAGCCGCTTGCT | GGG |
| Dux_KO_B | GACTTTCCCCACTAGTGGCT | TGG |
| Obox4_KO_8 | GGAAGAGCTGTACCCCTAGA | GGG |
| Obox4_KO_10 | GAGAAGGAACAATGTGCCTA | TGG |
